# *Disrupted in Renal Carcinoma 3* (*DIRC3*) impacts malignant phenotype and IGFBP5/IGF-1/Akt signaling axis in differentiated thyroid cancer

**DOI:** 10.1101/2023.01.24.525402

**Authors:** Piotr T. Wysocki, Karol Czubak, Anna A. Marusiak, Monika Kolanowska, Dominika Nowis

**Affiliations:** Laboratory of Experimental Medicine, Medical University of Warsaw, Nielubowicza 5, 02-097 Warsaw, Poland; Department of Oncology, Medical University of Warsaw, Banacha 1A, 02-097 Warsaw, Poland; Laboratory of Molecular OncoSignalling, IMol Polish Academy of Sciences, Flisa 6, 02-247 Warsaw, Poland; Warsaw Genomics INC, Łowicka 35, 02-502 Warsaw, Poland; Department of Immunology, Medical University of Warsaw, Nielubowicza 5, 02-097 Warsaw, Poland

**Keywords:** *DIRC3*, thyroid cancer, *IGFBP5*, IGF-1, invasiveness

## Abstract

Differentiated thyroid cancers (DTCs) are malignancies with ill-defined hereditary predisposition. Some germline variants influencing the risk of DTCs localize in *disrupted in renal carcinoma 3* (*DIRC3*), a poorly characterized long non-coding RNA (lncRNA) gene. Here, we characterized the function of *DIRC3* in DTCs. We established that *DIRC3* is downregulated in DTCs, and its high expression may reduce the risk of cancer recurrence in patients. *DIRC3* transcripts were enriched in cell nuclei *in vitro*, where they upregulated *insulin-like growth factor binding protein 5* (*IGFBP5*), a gene known to modulate the cellular response to insulin-like growth factor 1 (IGF-1). Silencing of *DIRC3* in thyroid cancer cell lines produced a phenotypic dichotomy: it augmented cell migration and invasiveness, reduced apoptosis, but abrogated the MTT reduction rate. We demonstrated that the pro-migratory phenotype was produced by the downregulation of *IGFBP5*. Transcriptomic profiling confirmed a functional redundancy in the activities of *DIRC3* and *IGFBP5*. Moreover, downregulation of *DIRC3* enhanced the susceptibility of cancer cells to IGF-1 stimulation and promoted Akt signaling. In conclusion, *DIRC3* expression alters the phenotype of thyroid cancer cells and modulates the activity of IGFBP5/IGF-1/Akt axis. We propose an interplay between *DIRC3* and IGF signaling as a mechanism that promotes thyroid carcinogenesis.

## INTRODUCTION

Thyroid cancer is the most common malignancy of endocrine glands.^1^ More than 95% of thyroid cancers originate from follicular epithelial cells.^2^ The current 4th World Health Organization Classification of Tumours of Endocrine Organs divides follicular cell-derived thyroid cancers into five entities: papillary (PTC), follicular (FTC), Hürthle cell (HTC), poorly differentiated, and anaplastic carcinomas. PTC, FTC and HTC retain a significant degree of cytological follicular differentiation and have a favorable prognosis. Accordingly, these three cancer types are jointly termed “differentiated thyroid cancer” (DTC).^2^

Contribution of hereditary factors to the pathogenesis of DTCs is one of the highest among all cancer types.^3, 4^ Pedigree studies demonstrate three- to nine-fold increase in thyroid cancer risk in the first-degree relatives of DTC patients.^3-5^ Furthermore, a recent pan-cancer study has indicated that thyroid cancers have the second highest heritability estimate among 18 common malignancies.^6^ Some insights into the mechanism of this hereditary predisposition have been provided by genome-wide association studies (GWAS).

At least seven GWAS have been performed in DTCs to date.^7-11^ These efforts have identified at least 10 chromosomal loci modulating the risk of thyroid cancer in the Caucasians. Four loci (14q13.3, 9q22.33, 8q12 and 2q35) contain germline variants demonstrating particularly robust associations and good cross-study replicability in European, American and Asian populations.^7-11^ Numerous single nucleotide polymorphisms (SNPs) associated with the risk of thyroid cancers locate in the chromosome 2q35 in *disrupted in renal carcinoma 3* (*DIRC3*), a poorly characterized long non-coding RNA (lncRNA) gene. *DIRC3* variants present some of the strongest associations with DTC incidence across ethnically diverse populations.^7-9, 12-14^ Additionally, one of the SNPs in *DIRC3*, rs996423, has been shown to influence the overall mortality in DTC patients.^15^

While the associations between *DIRC3* germline variants and the thyroid cancer risk have been robustly documented, the function of *DIRC3* in DTCs has not been established so far. Henceforth, we evaluated the role of *DIRC3* in the clinical and phenotypic presentation of DTCs.

## RESULTS

### *DIRC3* expression in public RNA-seq datasets

Evaluation of publicly available RNA-sequencing datasets (the Genotype-Tissue Expression data [GTEx] for normal tissue, and The Cancer Genome Atlas [TCGA] for cancers^16, 17^) revealed that *DIRC3* is expressed in normal thyroid tissue and PTCs (Figure 1A and Supp. Figure 1). Using a best fitting threshold of *DIRC3* expression to discriminate the disease-free status, we classified PTCs as either *DIRC3-high* or *DIRC3-low* tumors. Survival analysis indicated that elevated expression of *DIRC3* associated with a significantly longer disease free-survival (Figure 1B and 1C). A similar trend was observed for the overall survival (Supp. Figure 2). Histological make-up of *DIRC3-high* and *DIRC3-low* groups was dissimilar: *DIRC3-high* carcinomas were enriched for conventional PTCs (81.9% and 67.3% in *DIRC3-high* and *DIRC3-low* groups, respectively), but depleted of the follicular (12.6% and 23.1%, respectively) and tall-cell PTC variants (2.4% and 8.9%, respectively). Likewise, the prevalence of tumor-driving mutations was divergent in the expression groups (Supp. Figure 3).

**Figure 1.**
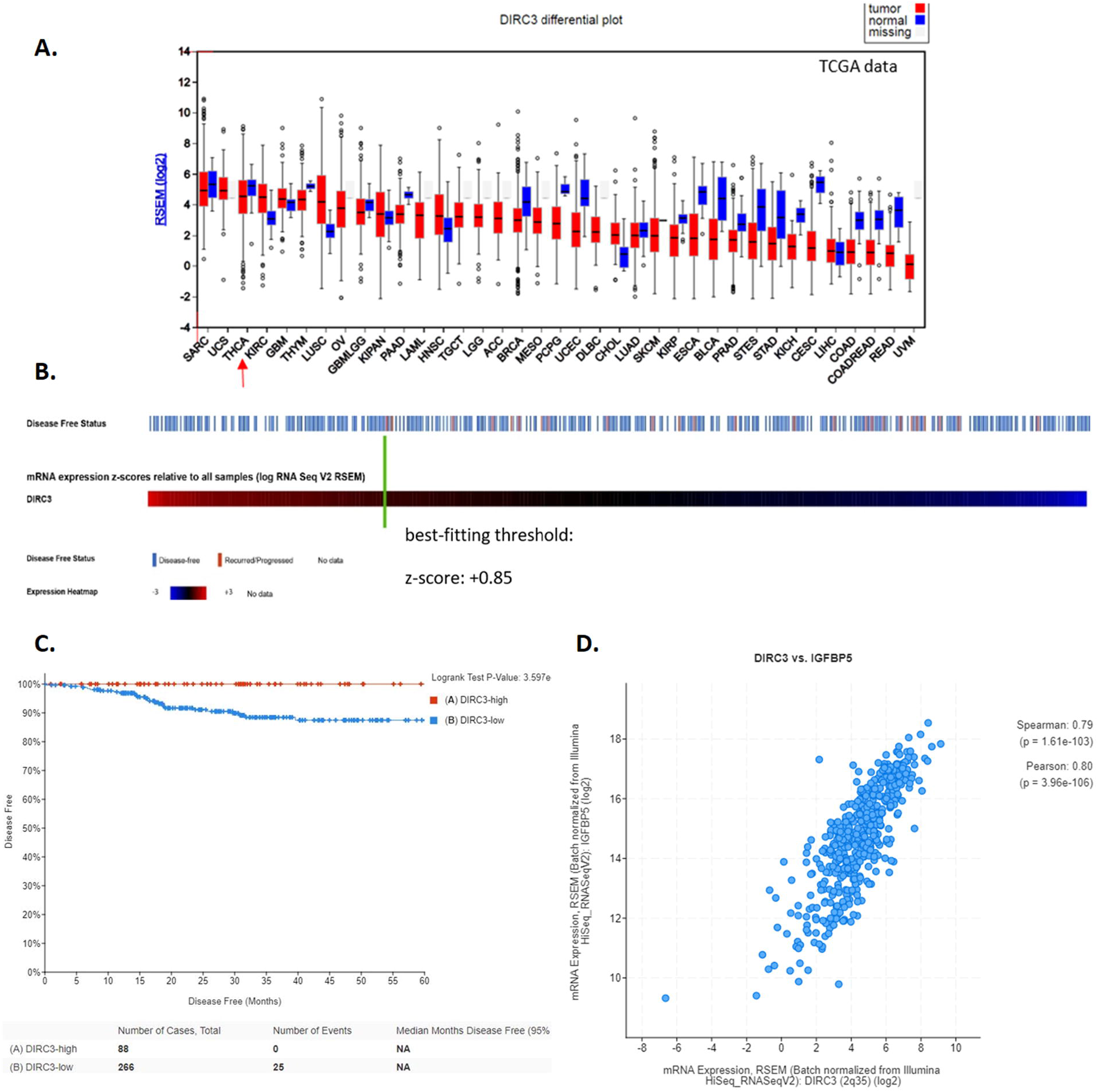
*DIRC3* expression in PTCs in TCGA. **(A)**. Expression of *DIRC3* in malignancies profiled in TCGA project. Data for PTCs (THCA) is marked with a red arrow (graph generated in the Firehose portal, http://firebrowse.org/, accessed 05/2018). **(B)**. Oncoprint graph illustrating the relationship between *DIRC3* expression and the disease-free status in PTCs. The best fitting threshold value of gene expression used to discriminate the disease-free status is marked with a green line. **(C)**. Disease-free survival of PTC patients according to the *DIRC3* expression status. **(D)**. Correlation between the expression of *DIRC3* and *IGFBP5* in PTCs in TCGA. Graphs were prepared using cBioPortal (https://www.cbioportal.org/; 05/2018). Abbreviation: RSEM, RNA-Seq by Expectation-Maximization.

TCGA data was used to evaluate correlations between the level of *DIRC3* lncRNA and expression of other genes (Supp. Table 1). *DIRC3* was most strongly co-expressed with *insulin-like growth factor binding protein 5, IGFBP5* (Spearman coefficient = 0.79), a gene located on the chromosome 2 approximately 570 000 base pairs downstream from *DIRC3* (Figure 1D).

### *DIRC3* expression in thyroid tissue

We analyzed the expression of *DIRC3* and *IGFBP5* in 67 DTC and normal thyroid tissue pairs (clinical data shown in Supp. Table 2). *DIRC3* was significantly downregulated in DTCs compared to the patient-matched normal thyroid tissue specimens (Figure 2A). Moreover, the strong co-expression of *DIRC3* and *IGFBP5* was confirmed (Figure 2B). This outcome suggested that *DIRC3* could play some role in the transcriptomic regulation of *IGFBP5*, a gene prominently involved in the modulation of insulin-like growth factor (IGF) signaling.

**Figure 2.**
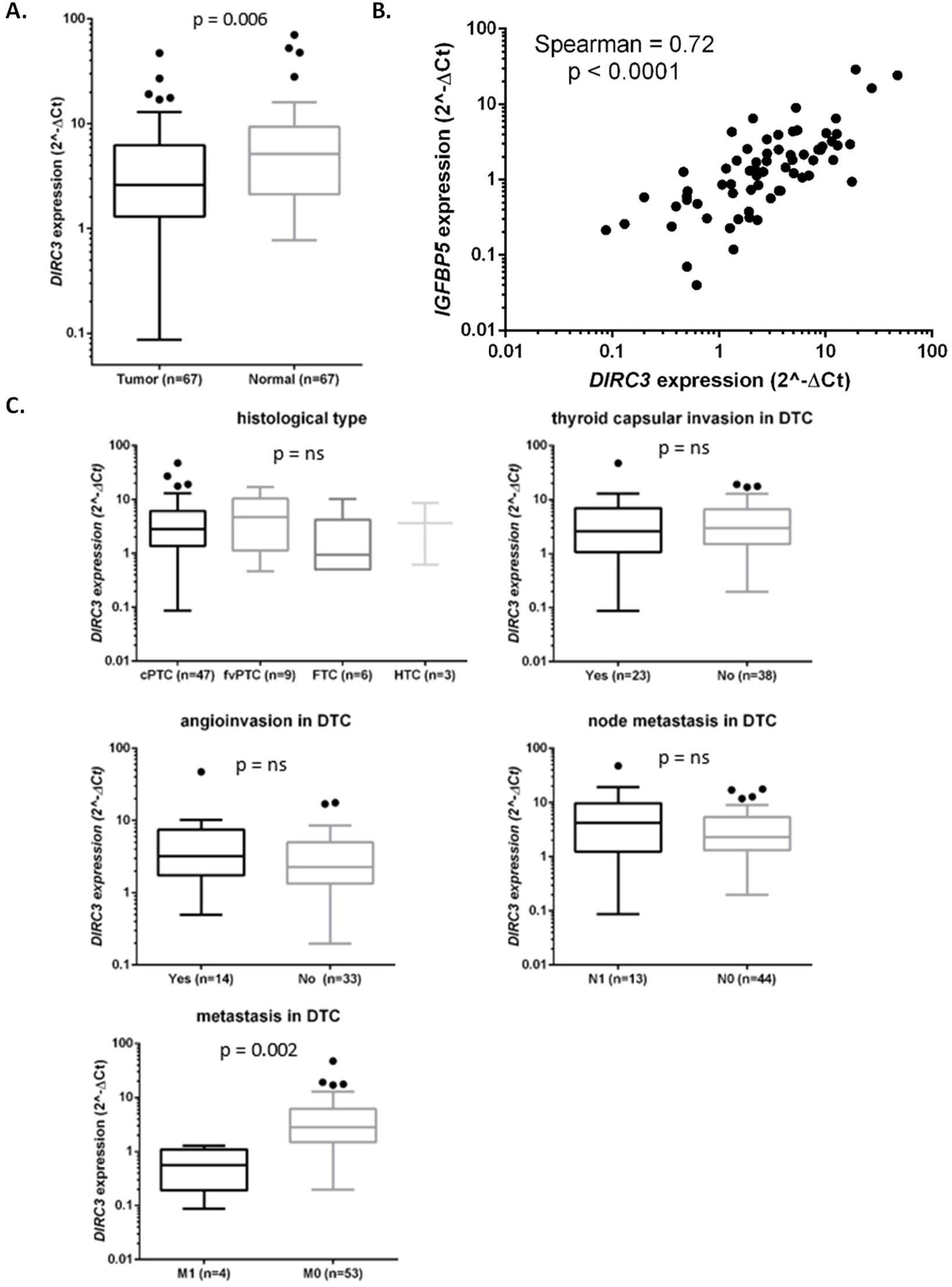
*DIRC3* expression in thyroid tissue. **(A)**. Expression of *DIRC3* in DTCs and normal thyroid tissue (Tukey plot; pair-matched Wilcoxon test). **(B)**. Correlation between the expression of *DIRC3* and *IGFBP5* in DTCs. Gene co-expression in normal thyroid tissue (Spearman coefficient = 0.77, p < 0.001) is not shown. **(C)**. Relationship between *DIRC3* expression in cancer and the clinicopathological features of DTCs (analyzed with Kruskal-Wallis and Mann–Whitney U tests). Abbreviations: cPTC: conventional PTC; fvPTC follicular variant PTC; FTC: follicular thyroid cancer; HTC: Hürthle cell thyroid carcinoma.

The expression of *DIRC3* was analyzed in the context of patients’ clinicopathological data (Figure 2C). Histological cancer type, invasion of thyroid capsule, node metastasis, or vascular invasion did not associate with the level of *DIRC3* lncRNA. On the other hand, *DIRC3* expression was lower in primary DTCs that developed distant metastasis. Separate analysis performed for conventional PTCs (cPTCs) did not identify significant interactions between *DIRC3* expression and clinicopathological features (Supp. Figure 4). No separate evaluations were attainable for other histological types due to their limited representation in our material.

### *DIRC3* expression in cancer cell lines

GTEx data revealed that *DIRC3* has four main splice variants (Supp. Figure 5). Only two splice variants are prominently expressed across all tissue types evaluated in GTEx. The longer splice variant is annotated as ENST00000474063.5 (*DIRC3-202* in Ensebl), and the shorter splice variant is ENST00000484635.1 (*DIRC3-203)*. Two other annotated splice variants do not exhibit expression in thyroid in the GTEx data: ENST00000486365.5 (*DIRC3-204*) and ENST00000423123.1 (*DIRC3-201*).

The expression of *DIRC3* and *IGFBP5* was evaluated in five cancer cell lines: K1, MDA-T32, MDA-T68, MDA-T120 (PTC cell lines) and MCF-7 (breast cancer cell line). We included MCF-7 due to previously reported associations of SNPs in *DIRC3* with the breast cancer risk^18, 19^, and a relatively high expression of *DIRC3* in our preliminary experiments. The highest expression of *DIRC3* and *IGFBP5* was observed in MDA-T32 and MCF-7, while *DIRC3* was not expressed in MDA-T68 (Figure 3A). We confirmed expression of *DIRC3-202* and *DIRC3-203* splice variants, and the absence of *DIRC3-204* in all tested cell lines. Subcellular fractionation of RNA showed that *DIRC3* transcripts localized preferentially in the nuclear fraction (Figure 3B).

**Figure 3.**
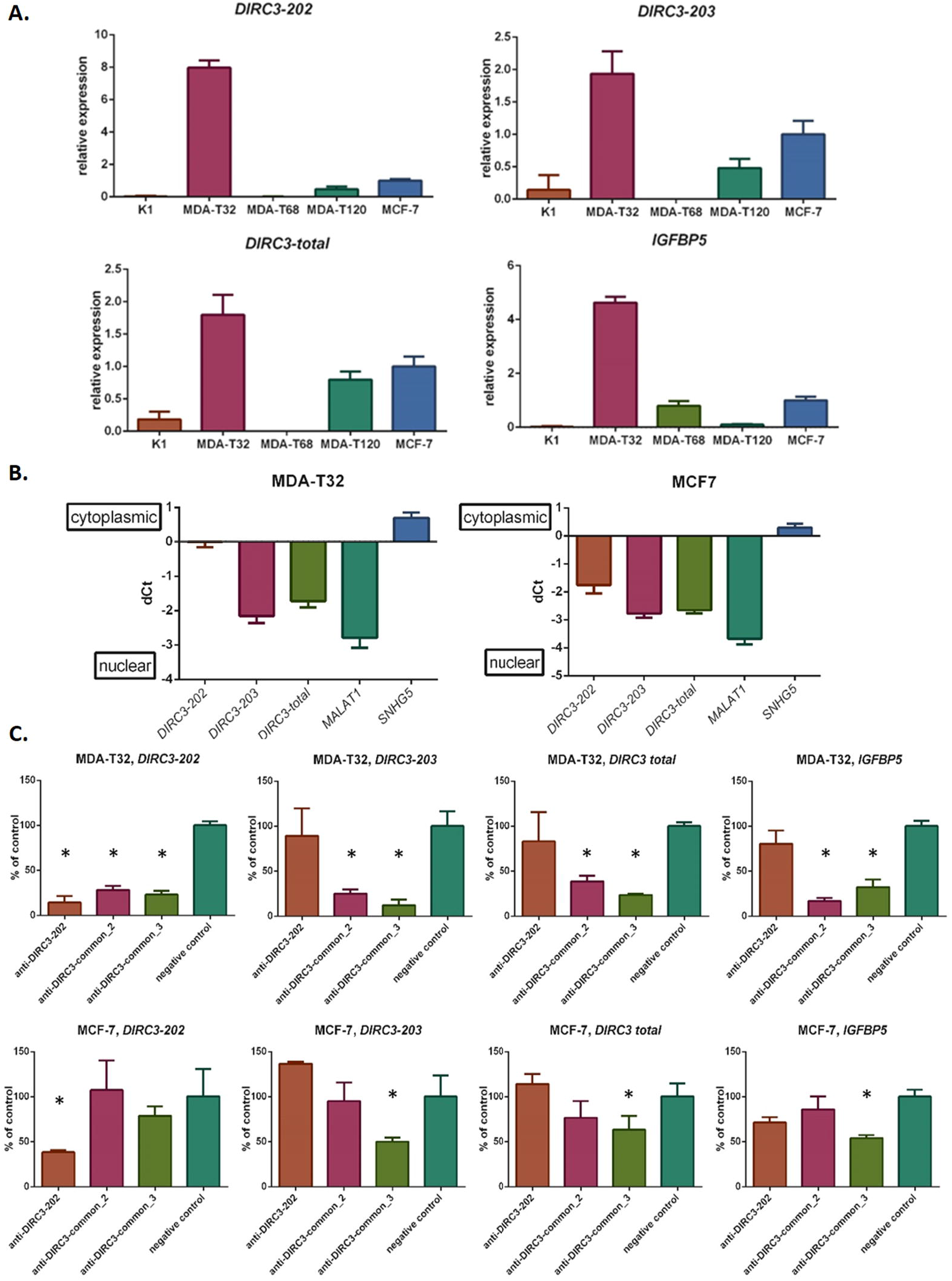
Expression and silencing of *DIRC3* in cancer cell lines. **(A)**. Gene expression (*DIRC3* total, *DIRC3* splice variants, and *IGFBP5*) in cell lines. *DIRC3-204* was not detected in any cell line (not shown). **(B)**. Subcellular localization of *DIRC3* and control (*MALAT1* and *SNHG5*) lncRNAs. Enrichment of transcripts in the subcellular fractions is expressed as ΔCt between the cytoplasmic and nuclear RNA fractions. **(C)**. Effect of *DIRC3* silencing in MDA-T32 and MCF-7 cells on the expression of *DIRC3* (total and splice variants) and *IGFBP5*. * indicates p<0.05 vs. negative control (n = 3; mean ± SD; ANOVA).

We used *DIRC3*-targeting antisense oligonucleotides to silence *DIRC3* expression in four cell lines (Figure 3C and Supp. Figure 6). Two GapmeRs (anti-*DIRC3*-common_2 and anti-*DIRC3*-common_3) were designed to silence both expressed splice variants (*DIRC3-202* and *DIRC3-203*). Additionally, another GapmeR (anti-*DIRC3-202*) targeting only the longer splice variant was used to test the phenotypic effects of selective silencing of this transcript. *DIRC3*-targeting GapmeRs successfully downregulated their direct targets. Importantly, silencing of *DIRC3* also downregulated *IGFBP5* in the cell lines that expressed *DIRC3* (MDA-T32, MDA-T120 and MCF-7). Interestingly, GapmeR anti*-DIRC3-202* successfully silenced *DIRC3-202*, however this downregulation did not influence *IGFBP5*. Finally, *DIRC3*-targeting GapmeRs did not influence *IGFBP5* expression in MDA-T68, the cell line not expressing *DIRC3* (Figure 3). This result proved that *IGFBP5* downregulation was directly related to the *DIRC3*-targeting capabilities of the GapmeRs.

### Phenotypic impact of *DIRC3* silencing

The phenotypic effects of *DIRC3* silencing were evaluated in MDA-T32 and MDA-T120 cell lines. We utilized MTT assays to indirectly quantify the cell proliferation and viability. Downregulation of *DIRC3* modestly restrained the MTT reduction rate in MDA-T32, while a prominent inhibitory effect was observed in MDA-T120 (Figure 4A). Interestingly, no such result was observed when *DIRC3-202* was silenced selectively. This outcome might indicate different biological and transcriptomic activities of *DIRC3* splice variants.

**Figure 4.**
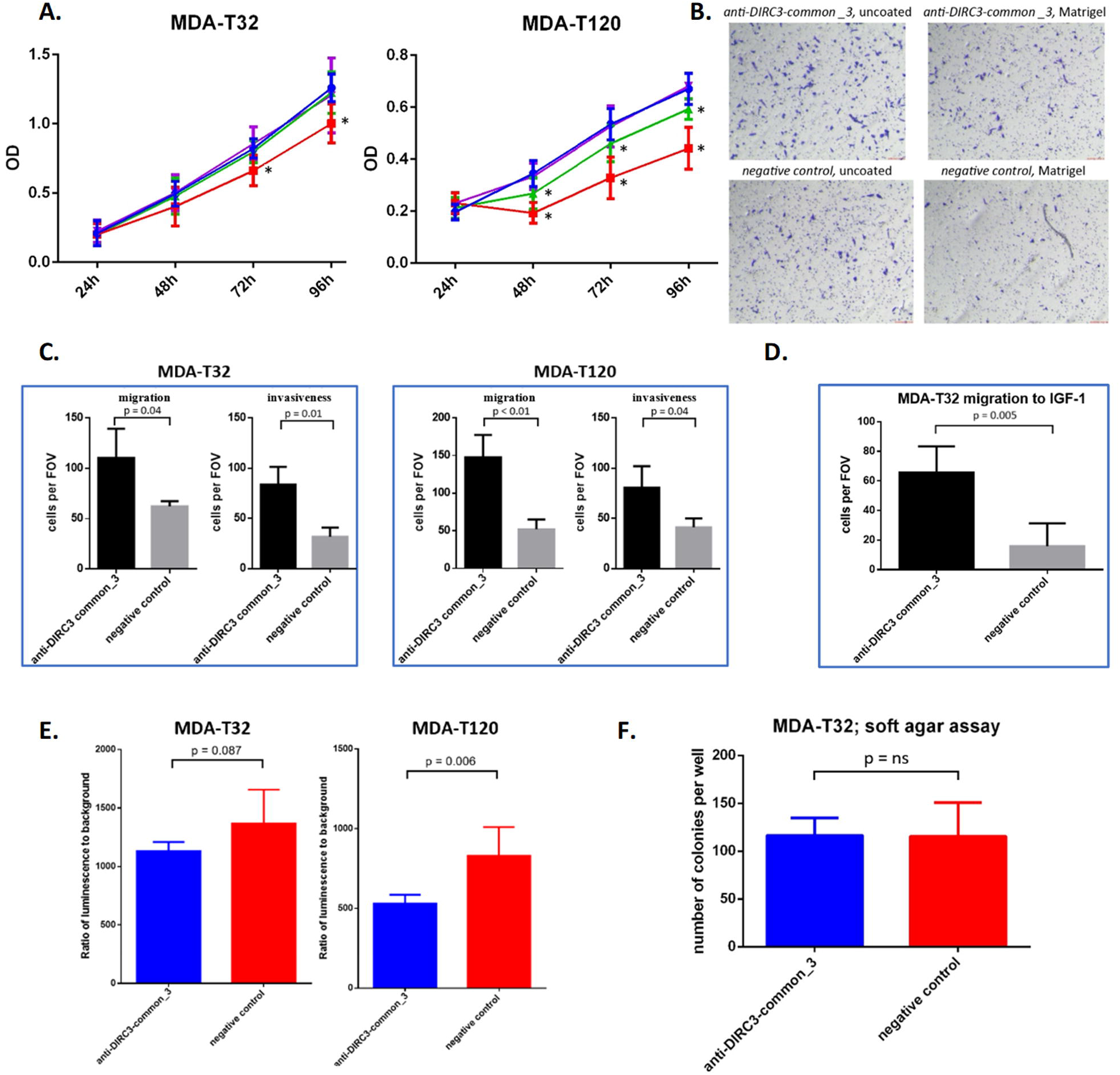
Phenotypic effect of *DIRC3* silencing in MDA-T32 and MDA-T120 cell lines. **(A)**.Results of MTT assays after GapmeR transfections. * indicates p < 0.05 vs. negative control (n = 3 with technical quaruplicates; mean ± SD; ANOVA). **(B)**. Representative images of the GapmeR-transfected MDA-T32 cells in Transwell assays. **(C)**. Quantification of migration and invasiveness in MDA-T32 and MDA-T120 cells after silencing of *DIRC3* (n = 3; mean ± SD; t-test). **(D)**. Quantification of the chemotactic effect of IGF-1 in *DIRC3*-depleted MDA-T32 cells (n = 3; mean ± SD; t-test). **(E)**. Activity of caspase 3/7 after GapmeR transfections in serum- and glutamine-starved MDA-T32 and MDA-T120 cells (n = 6; mean ± SD; t-test). **(F)**. Number of visible colonies per well in soft agar assays after transfection of MDA-T32 cell with the *DIRC3*-silencing and negative control GapmeRs (n = 9; mean ± SD; t-test). Abbreviations: FOV, field of view; ns: not significant; OD, optical density.

Downregulation of *DIRC3* markedly increased migration and invasiveness of MDA-T32 and MDA-T120 cell lines in the Transwell assays (Figure 4B & 4C). Similarly, silencing of *DIRC3* in MDA-T32 cells significantly promoted their chemotaxis to IGF-1 (Figure 4D). IGF-1 alone was insufficient to generate a chemotactic response in MDA-T120 (not shown).

We tested if the alterations in *DIRC3* expression impact the apoptosis susceptibility of cancer cells deprived of serum and L-glutamine. A luminescent assay indicated that silencing of *DIRC3* reduced the activity of caspase 3/7 in starved MDA-T120 and MDA-T32 cells (Figure 4E).

Silencing of *DIRC3* did not influence the anchorage-independent growth capabilities of MDA-T32 cells (Figure 4F and Supp. Figure 7). MDA-T120 cell line generated only a limited number of visible colonies in soft agar thus preventing its use in the assay.

### *IGFBP5* rescue experiments

We used rescue experiments involving *IGFBP5* to establish whether phenotypic effects induced by *DIRC3* silencing were related to the transcriptomic regulation of *IGFBP5*. For this purpose, two MDA-T32 derivatives with stable transgene expression were generated: cells expressing pcDNA3-IGFBP5-V5 plasmid, and cells transfected with an empty pcDNA3 vector (a negative control).

Both modified cell lines were transfected with the *DIRC3*-targeting GapmeR (Figure 5A). Introduction of pcDNA3-IGFBP5-V5 to MDA-T32 resulted in a strong upregulation of *IGFBP5*. Transfection of anti-*DIRC3*-common_3 downregulated *DIRC3* in both plasmid-overexpressing cell lines. Nevertheless, this procedure significantly downregulated *IGFBP5* only in the cells transfected with the control plasmid (Figure 5A). This outcome indicated that the *DIRC3*-targeting GapmeR could influence *IGFBP5* expression only when the transcripts were produced from the nuclear locus. Besides, pcDNA3-IGFBP5-V5 modestly upregulated the expression of *DIRC3*.

**Figure 5.**
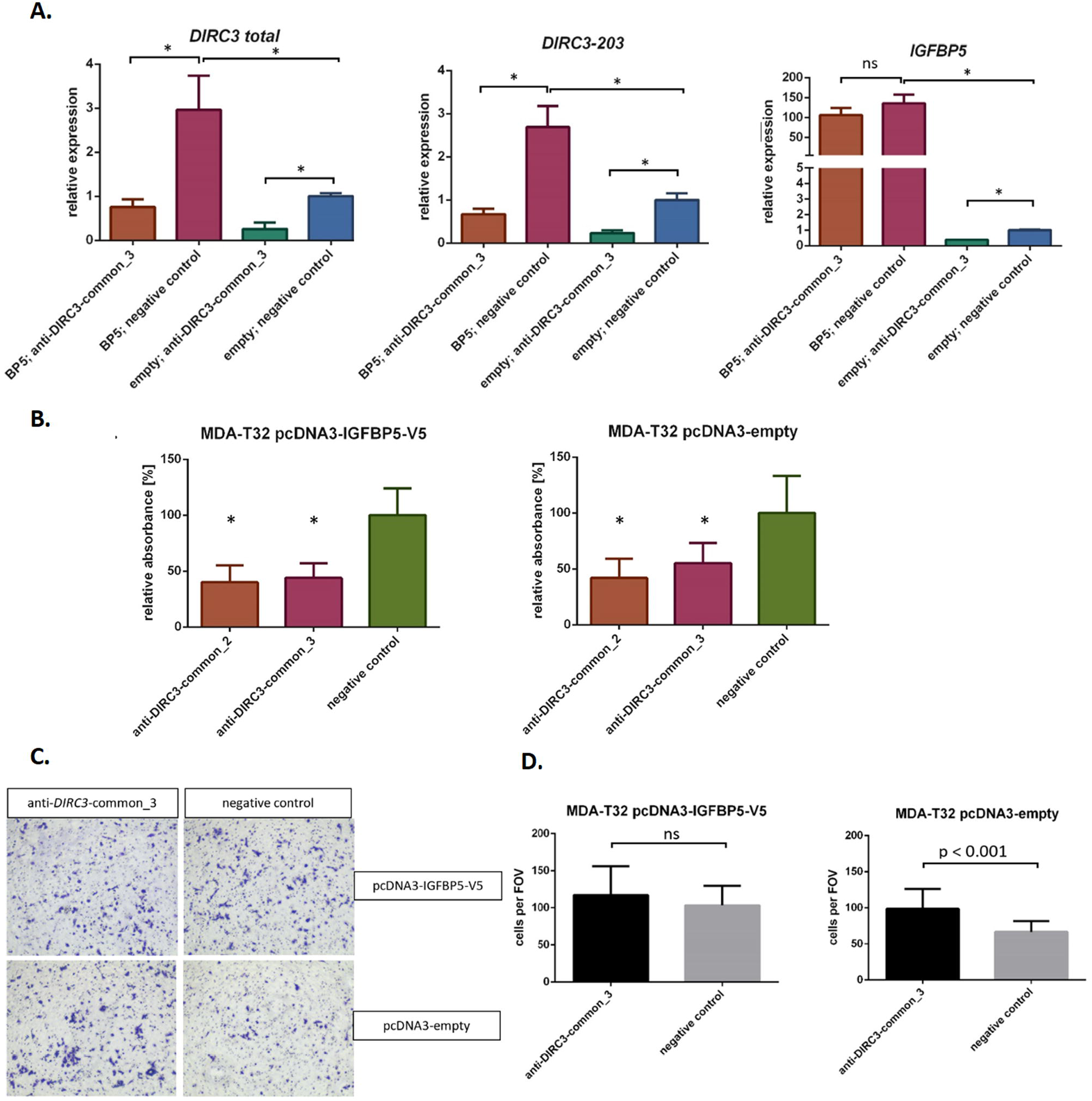
*IGFBP5*-rescue experiements in MDA-T32 cells transfected with the *DIRC3*-targeting GapmeRs. **(A)**. Expression of *DIRC3* (total and *DIRC3-203*) and *IGFBP5* in MDA-T32 cells with stable expression of pcDNA3-IGFBP5-V5 (abbreviated as “BP5”) or control (“empty”) pcDNA3 plasmids, transfected with GapmeRs (n = 3; mean ± SD; t-test). **(B)**. Results of MTT assays in the plasmid-expressing MDA-T32 cells 96 hours after transfections of GapmeRs (n = 3 with technical quadruplicates; mean ± SD; ANOVA vs. control). **(C)**. Representative fields of view (FOV) in Transwell assays of the plasmid-expressing and GapmeR-transfected MDA-T32 cells. **(D)**. Quantification of migration of the plasmid-expressing MDA-T32 cells after transfections of GapmeRs (n = 3; mean ± SD, t-test). * indicates p < 0.05 vs control. ns: not significant.

Plasmid-transfected MDA-T32 cells were tested in MTT and Transwell assays. *DIRC3* silencing significantly reduced the MTT conversion rate in both plasmid-expressing cell derivatives (Figure 5B). In contrast, overexpression of *IGFBP5* successfully negated the pro-migratory phenotype generated by *DIRC3* silencing (Figure 5C and 5D). These results suggested that the phenotypic effects produced by *DIRC3* could be either *IGFBP5*-dependent (in regards to the migratory potential), or at least partially independent from *IGFBP5* (as in MTT assays).

### Transcriptomic alterations induced by *DIRC3* silencing

We used RNA-seq to evaluate transcriptomic changes related to the silencing of *DIRC3* or *IGFBP5* in MDA-T32. Efficient gene downregulation was confirmed using qRT-PCR (Supp. Figure 8). As expected, *DIRC3* silencing reduced the abundance of *IGFBP5* transcripts. Interestingly, the knockdown of *IGFBP5* also downregulated *DIRC3*. This observation was in line with the outcomes of *IGFBP5* overexpression experiment, in which *IGFBP5* upregulated *DIRC3*. Accordingly, a bidirectional positive feedback mechanism between *DIRC3* and *IGFBP5* may be proposed.

RNA-seq was performed in biological triplicates for each set-up (the primary component analysis shown in Supp. Figure 9). Silencing of *DIRC3* revealed 198 differentially expressed genes (DEGs), while silencing of *IGFBP5* resulted in 631 DEGs. Heatmaps of top 30 DEGs in *DIRC3*-silenced and *IGFBP5*-silenced groups, and corresponding volcano plots are shown in Figure 6A and Supp. Figure 10, respectively. Gene overlap between the *DIRC3*- and *IGFBP5-* regulated DEGs was significant and comprised of 58 genes (Figure 6B). Directions of the expression changes were concordant for all shared DEGs. Review of literature revealed that many of the shared DEGs have been previously implicated in thyroid carcinogenesis, e.g., *metastasis associated lung adenocarcinoma transcript 1* (*MALAT1), matrix metalloproteinase-1* (*MMP-1*), *cyclin dependent kinase inhibitor 1A* (*CDKN1A*) and *stanniocalcin 1* (*STC1*) (Supp. Table 3).

**Figure 6.**
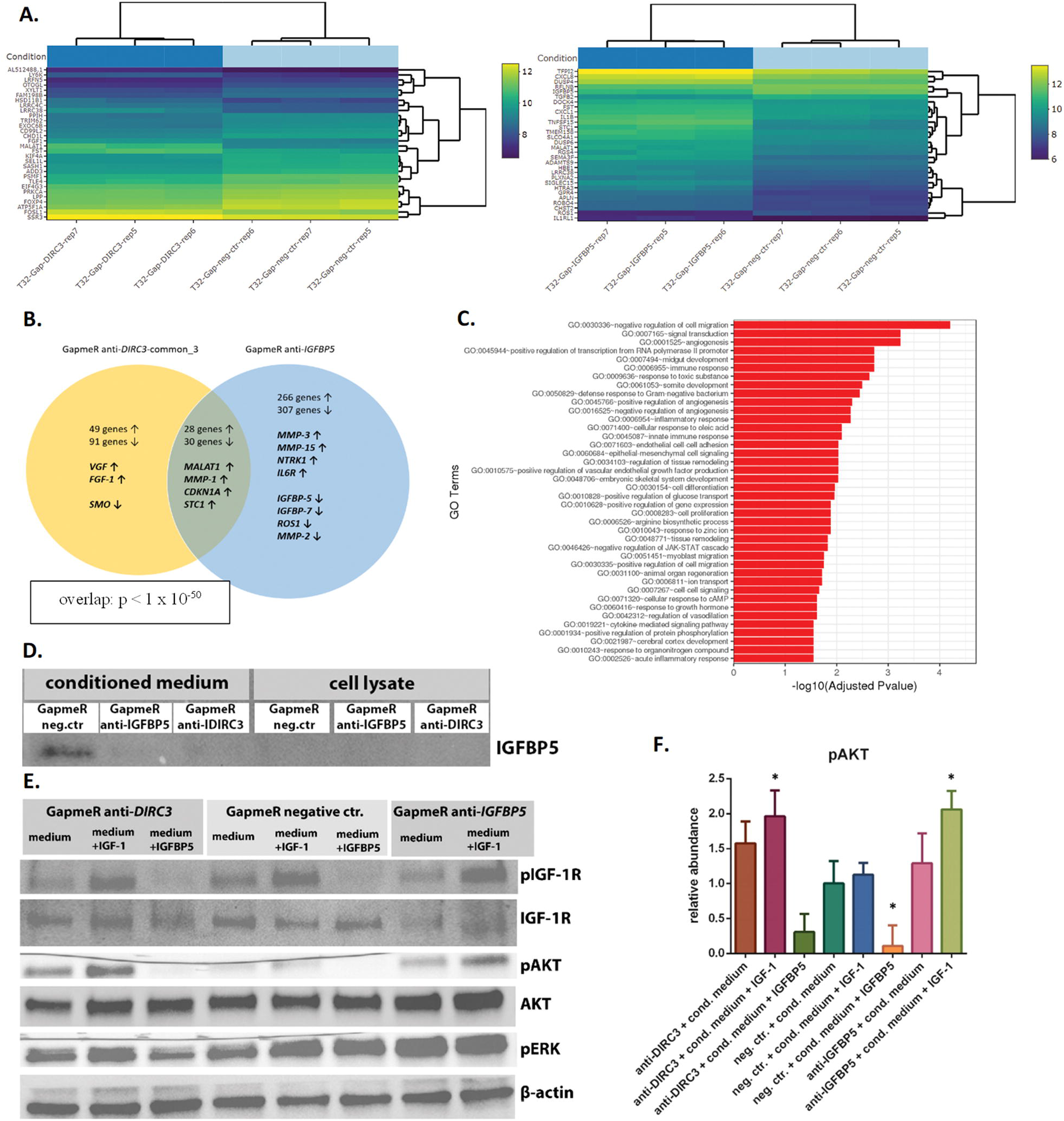
Influence of *DIRC3* and *IGFBP5* downregulations on the transcriptome and IGF-1R/AKT signaling in MDA-T32 cells. **(A)**. Bi-clustering heatmaps of top 30 *DIRC3*- and *IGFBP5*-altered DEGs (log2 transformed values; sorted by adjusted p-values). **(B)**. Overlap between DEGs in the *DIRC3*- and *IGFBP5*-silenced samples (n = 3). Arrows indicate up- or down-regulation. **(C)**. Top gene ontology (GO) terms enriched in MDA-T32 cells after silencing of *DIRC3*. **(D)**. Immunoblot of IGFBP5 in the conditioned media and MDA-T32 cells tranfected with GapmeRs (cultured with 20 ng/ml of IGF-1, 24 h). **(E)**. Immunoblots of MDA-T32 cells transfected with GapmeRs and stimulated for 10 mins with previously harvested conditioned media. The conditioned media were applied in 3 versions: a) unmodified (i.e. with IGF-1 20 ng/ml), b) supplemented with additional 30 ng/ml of IGF-1, or c) supplemented with IGFBP5 (500 ng/ml). **(F)**. Relative abundance of pAKT in three independent blots (mean ± SD of semi-quantitative densitometric units; ANOVA). * indicates p < 0.05 vs. “neg. ctr. + cond. medium”.

The gene ontology (GO) analysis indicated that genes involved in the “negative regulation of cell migration” (GO:0030336) were most significantly affected by the knockdown of *DIRC3* (Figure 6C). Other GO terms significantly altered included: “signal transduction” (GO:0007165), “cell proliferation” (GO:0008283), “response to growth hormone” (GO:0060416), “positive regulation of protein phosphorylation” (GO:0001934). *IGFBP5* is assigned to 29 GO terms in The Gene Ontology Annotation Database. Several of these terms were among the most significantly altered by *DIRC3* silencing (e.g., GO:0030336∼negative regulation of cell migration, GO:0007165∼signal transduction, GO:0071320∼cellular response to cAMP).

### Effect of *DIRC3* silencing on IGF signaling

Since *DIRC3* modulated the expression of *IGFBP5*, we hypothesized that *DIRC3* could impact IGF signaling. Silencing of *DIRC3* or *IGFBP5* in MDA-T32 cells downregulated the release of IGFBP5 into medium containing IGF-1 (Figure 6D). We harvested the conditioned media and starved cells for 12 h. Stimulation of the starved cells with their original conditioned medium induced phosphorylation of IGF-1 receptor (IGF-1R) and Protein kinase B (Akt; Figure 6E and 6F). The positive influence on pAkt level was significantly stronger in the cells transfected with either *DIRC3*- or *IGFBP5*-targeting GapmeRs (i.e. when the conditioned medium contained less IGFBP5). Phosphorylation of Akt was further increased when the conditioned media was enhanced with additional IGF-1. On the other hand, supplementation of recombinant IGFBP5 protein prevented phosphorylation of IGF-1R and Akt. These outcomes indicated that the strong phosphorylation of Akt observed after silencing of *DIRC3* was triggered by the stimulatory effect of IGF-1 and reduced amount of IGFBP5 in the medium. In contrast, extracellular signal-regulated kinase (ERK) was robustly phosphorylated in all samples since MDA-T32 harbors *BRAF* V600E mutation (typical for cPTC). In conclusion, downregulation of *DIRC3* influenced response to IGF-1 and promoted Akt signaling in thyroid cancer cells. A hypothetical mechanistic model is shown in Supp. Figure 11.

## DISCUSSION

This study is the first to report that *DIRC3* is functionally implicated in thyroid carcinogenesis. We demonstrate that *DIRC3* is downregulated in DTCs, and its low expression may increase the risk of cancer recurrence. Silencing of *DIRC3* in thyroid cancer cells augmented their invasiveness, curtailed production of IGFBP5 and boosted Akt signaling upon IGF-1 stimulation. Transcriptomic profiling performed in cells experiencing knockdown of either *DIRC3* or *IGFBP5* indicated a significant redundancy in the activities of both genes. Shared and upregulated DEGs included many genes previously implicated in thyroid tumorigenesis, e.g., *MALAT1, CDKN1A* and *MMP-1*.^20-22^ *DIRC3* silencing also upregulated *STC1*, which encodes stanniocalcin 1, a potent inhibitor of pregnancy-associated plasma protein -A and -A2 (PAPP-A and PAPP-A2).^23^ PAPP-A and PAPPA-2 possess proteolytic activities towards IGFBP5. Stanniocalcin 1 thus prevents the release of IGF-1 from IGF/IGFBP complexes and downregulates IGF signaling.^23^ Upregulation of *STC1* may constitute a negative feedback mechanism triggered by the knockdown of *DIRC3* or *IGFBP5. STC1* also acts as an oncogene promoting proliferation of thyroid cancer cells.^23, 24^

*DIRC3* was originally described as a gene participating in t(2;3)(q35;q21) translocation in familial renal cell cancers, however, its function was not evaluated in the report.^25^ Critical evidence implicating *DIRC3* in carcinogenesis has been provided by GWAS. Germline variants in *DIRC3* were found to influence the hereditary risk of estrogen receptor-positive breast cancers.^6, 18, 19^ Certain breast cancer risk variants located in the chromosome 2q35 (in the proximity or within *DIRC3* locus) were mapped to putative regulatory elements (PREs) bound by estrogen receptor (ERα) and forkhead box A1 (FOXA1) transcription factors. Some of these PREs were found to act as enhancers that physically interact with the *IGFBP5* locus to regulate its expression.^26, 27^

Numerous germline variants in *DIRC3* have been associated with the risk of DTCs (rs966423, rs6759952, rs12990503, rs16857609, rs11693806 and rs772695095).^6, 8, 9, 13, 28, 29^ Guibon *et al*. have recently performed fine-mapping analysis of the *DIRC3* locus.^14^ This project replicated some of the previously reported risk SNPs and identified two novel risk variants (rs57481445 and rs3821098). Colocalization analysis highlighted three additional SNPs in *DIRC3*, which were found to constitute *expression quantitative trait loci* associated with the downregulation of *DIRC3* and *IGFBP5* in thyroid tissue. The strongest effect size was observed for rs12990503, the variant previously associated with increased risk of thyroid and breast cancers.^14, 29^

Coe *et al*. have recently demonstrated that two transcription factors critical in melanomagenesis, melanocyte inducing transcription factor (MITF) and SRY-related HMG-box 10 (SOX10), colocalize to two PREs in the *DIRC3* locus and suppress *DIRC3* expression.^30^ Since *DIRC3* and *IGFBP5* were located in a common topological domain, the authors hypothesized that both genes could be transcriptomically related. Indeed, silencing of *DIRC3* downregulated *IGFBP5* expression in melanoma cell lines and promoted their anchorage-independent growth. While the results of the melanoma study support our discoveries, it is important to stress differences.

Firstly, while Coe *et al*. observed that suppression of *DIRC3* promoted the anchorage-independent growth, no changes in the proliferation and migration of melanoma cells were detected.^30, 31^ This outcome contrasts our observations. Additionally, overexpression of *IGFBP5* in melanoma was previously shown to inhibit cell proliferation, anchorage-independent growth, migration and invasiveness *in vitro*, and reduce the melanoma growth and pulmonary metastasis *in vivo*. Conversely, silencing of *IGFBP5* in melanoma produced tumor-promoting effects.^32^ Secondly, the mechanistic model offered in melanoma may not hold true for thyroid cancers, since SOX10 and MITF are not expressed in normal thyroid and DTCs (as indicated by the GTEx and TCGA data). Thirdly, SNPs modulating the incidence of DTCs do not overlap PREs reported in the melanoma study. This suggests that these variants are very unlikely to impact functions of SOX10 and MITF. Fourthly, we observed that selective silencing of *DIRC3-202* did not influence *IGFBP5* expression. Interestingly, the sequence of our GapmeR anti-*DIRC3-202* was almost identical to the sequence of a GapmeRs used in the melanoma study. This particular GapmeR efficiently downregulated *IGFBP5* in melanoma cell lines. We presume that this inconsistency might indicate dissimilarities in the functions of *DIRC3* splice variants across various malignancies.

IGF signaling is strongly involved in carcinogenesis due to its pro-mitogenic, pro-invasive and anti-apoptotic roles. Its tumor-driving impact has also been recognized in DTCs.^33, 34^ The prospective UK Biobank study of 30 cancer types has recently reported that the concentration of IGF-1 in human serum was most strongly positively associated with the incidence of DTCs.^35^ Interestingly, several *DIRC3* SNPs associate with human height, the anthropometric parameter driven directly by IGF-1.^36, 37^ Given the strong positive association between body height and thyroid cancer risk, a common molecular mechanism driving both conditions may be presumed.^38, 39^ We propose that the epidemiological data associating human height with the incidence of DTCs can be at least partially explained by shared hereditary variants in *DIRC3* and their modulatory influence on the cellular response to IGF-1.

In the classical paradigm, IGFBP5 inhibits IGF signaling by preventing binding of IGF-1 to IGF-1R.^40^ Nevertheless, some studies demonstrate that IGFBP5 may promote IGF signaling in specific cellular and tissue contexts. The ultimate phenotypic influence of IGFBP5 is dependent on the bioavailability of IGF-1, abundance of IGFBP-degrading proteases, and the composition of extracellular matrix.^40^ Accordingly, *IGFBP5* may produce diverse biological effects. *IGFBP5* has been reported to act as either oncogene or tumor suppressor in different cancer studies, with some evidence indicating its pro-proliferative role in PTC.^40-42^ Results of our study designate *DIRC3* as functionally dichotomous gene, with its silencing boosting migration and invasiveness of cancer cells, but decreasing the MTT conversion rate (the indirect indicator of cell proliferation). We show that these phenotypic alterations may be either *IGFBP5*-dependent (changes in the migratory potential) or *IGFBP5*-independent (the effects observed in MTT assays). Accordingly, this functional dichotomy of *DIRC3* may echo the multifaceted phenotypic influence of *IGFBP5*, as well as correspond to some hypothetical *IGFBP5*-independent activities.

Our study has some limitations. Since the number of analyzed DTC samples was relatively small, our clinicial observations should be validated. Additionally, it is possible that our discoveries may hold true only for certain histological types of DTCs, principally PTCs. Moreover, we were unable to perform successful *DIRC3* overexpression experiments. While CRISPR activation (CRISPRa) was expected to recapitulate the endogenous expression of *DIRC3* more faithfully than the plasmid-mediated overexpression (which failed to produce biological effects in melanoma),^30^ our CRISPRa experiments were unsuccessful in upregulating *DIRC3* and *IGFBP5*. Future studies should also evaluate the role of *DIRC3 in vivo*.

Our discoveries may be clinically relevant. Firstly, expression of *DIRC3* in thyroid carcinomas may have a prognostic value, and the analysis of *DIRC3* level in thyroid tumors might help to guide medical decision (e.g., to deescalate treatment of low-risk/*DIRC3-high* cancers). Secondly, we demonstrate that downregulation of *DIRC3* increases the cellular response to IGF-1. Therapeutic inhibition of IGF signaling has been long pursued in clinical oncology.^43, 44^ While results of early studies of IGF-1R inhibitors were promising (with some trials enrolling DTC patients), outcomes of phase II/III clinical trials have been largely disappointing. Still, some exceptional responders were observed. It has been proposed that novel predictive factors are necessary to identify patients who could respond to IGF-1R inhibitors.^43, 44^ Remarkably, the level of IGFBP5 in tumors correlated inversely with the resistance to IGF-1R inhibition in breast, colon and bladder carcinomas.^45-47^ Accordingly, our results may provide rationale for evaluating IGF-1R inhibitors in DTCs that downregulate *DIRC3*.

In summary, our study indicates that *DIRC3* has a prominent anti-invasive role in thyroid cancers. *DIRC3* modulates the expression of *IGFBP5*, and hence it regulates IGF-1/Akt signaling. The expression of *DIRC3* in thyroid cancers may emerge as a clinically relevant prognostic factor and a predictive marker for novel therapeutic strategies.

## MATERIALS AND METHODS

### Clinical material

67 patient-matched DTC and normal thyroid tissue pairs were collected at the Medical University of Warsaw (Poland). Collection of tissue was approved by the Institutional Review Board. Informed consents were obtained from patients. RNA was extracted from fresh frozen tissue using TRIzol (Thermo Fisher, Waltham, MA, USA).

### Quantitative reverse transcriptase real-time PCR (qRT-PCR)

Reverse transcription was performed using the Moloney Murine Leukemia Virus Reverse Transcriptase (Promega, Madison, WI, USA), 1000 ng of total RNA and random hexamers. LightCycler 480 SYBR Green I Master Mix (Roche, Mannheim, Germany) was combined with diluted cDNA, and forward and reverse primers (0.4 μM final, each; Supp. Table 4). Reactions were performed in technical triplicates in LightCycler 480 II system (Roche). Relative gene expression was calculated using the 2^-∆∆Ct^ method employing *hypoxanthine phosphoribosyltransferase 1* (*HPRT1)* as the housekeeping gene.

### Bioinformatics databases

Gene expression was evaluated in public RNA sequencing data from: the Genotype-Tissue Expression project for normal tissue (https://gtexportal.org/home/), and The Cancer Genome Atlas for malignant tissue (accessed via cBioPortal; https://www.cbioportal.org/).

### Cell lines

MDA-T32 (RRID:CVCL_W913), MDA-T120 (RRID:CVCL_QW85; both conventional PTC), MDA-T68 (RRID:CVCL_QW83; follicular variant of PTC), and MCF-7 (RRID:CVCL_0031; breast cancer) cell lines were obtained from American Type Culture Collection (Manassas, VA, USA). K1 (RRID:CVCL_2537; conventional PTC) was obtained from Sigma-Aldrich (St. Louis, MO, UK). MDA-T32, MDA-T68 and MDA-T120 cells were cultured in RPMI-1640 (ATCC modification; Gibco, Paisley, UK). K1 was cultured using a mix of DMEM, Ham′s F12 (both Gibco, Grand Island, NY, USA) and MCDB-105 (Cell Applications, San Diego, CA, USA; ratio 2:1:1). Culture media were supplemented with 10% fetal bovine serum (FBS; Euroclone, Pero, Italy), 2 mM L-glutamine (Lonza, Walkersville, MD, USA), Penicillin-Streptomycin (Sigma-Aldrich), and MEM NEAA solution (Gibco; added only to RPMI-1640). Cultures were periodically tested for mycoplasma. Total RNA was extracted using the GeneMATRIX Universal RNA Purification Kit (EURx, Gdansk, Poland).

### Subcellular RNA fractionation

RNA fractionation of MDA-T32 and MCF-7 cells was performed using the Nuclei EZ lysis buffer (Sigma-Aldrich) as previously described.^48^ RNA was isolated from the nuclear and cytoplasmic cell fractions using RNA Extracol and GeneMATRIX Universal RNA Purification Kit (EURx). qRT-PCR was performed for each RNA fraction. The relative subcellular location of transcripts was estimated by calculating differences in the cycle-threshold values obtained for equal input (1000 ng) of nuclear and cytoplasmic RNA. *MALAT1* and *small nucleolar RNA host gene 5* (*SNHG5)* were used as nuclear and cytoplasmic control lncRNAs, respectively. Experiments were repeated three times.

### Gene silencing

Cells were plated in 6-well plates 24 h before transfections (3 × 10^5^ cells/well for MDA-T32, MDA-T68, K1 and MCF-7; 4 × 10^5^ cells/well for MDA-T120). GapmeRs (Qiagen, Hilden, Germany; 50 nM) were transfected using Lipofectamine 2000 (Thermo Fisher). Sequences of GapmeRs are provided in Supp. Table 5. In MTT assays transfections were performed directly in 96-well plates.

### Migration and invasiveness

Cells were tested using 8.0 µm pore Transwell chambers placed in 24-well plates (uncoated or Matrigel-coated inserts; Corning, Bedford, MA, USA). Cells were cultured for 48 h after transfections, serum-starved for another 24 h, dissociated using the Non-enzymatic Cell Dissociation Solution (Sigma-Aldrich), centrifuged, and re-suspended in serum-free medium. 5 × 10^4^ viable cells were applied to the upper compartments of inserts, while the lower compartments were filled with complete culture medium (10% FBS). Inserts were fixed with 4% paraformaldehyde after 22 h of culture, stained with 0.05% crystal violet, and rinsed with water. Upper surfaces of the insert membranes were cleansed with cotton swabs. Membranes were visualized (40x magnification), and at least 5 random fields of view were photographed in each insert. The IGF-1 chemotaxis assays were performed identically, except the serum-starvation step was omitted, and the lower compartments of chambers were filled with serum-free RPMI-1640 medium containing IGF-1 (100 ng/ml; Gibco). Each experiment was repeated at least three times.

### MTT assay

Cells were plated in 96-well plates (3 300 and 5 000 cells/well for MDA-T32 and MDA-T120, respectively) and transfected with GapmeRs. Culture medium was replaced with 100 μl OptiMEM (Gibco) immediately before testing. 10 μl of MTT solution (12 mM; Sigma-Aldrich) was added to the wells and incubated for 4 h. 100 µL of dissolving solution (0.1 g/ml SDS in 0.01 M HCl; Thermo Fisher, Rockfold, IL, USA) was added to each well. The optical density was measured spectrophotometrically at 560 nm (Glomax microplate reader, Promega) after 10 h of incubation. Experiments were performed with technical quadruplicates and repeated three times.

### Apoptosis assay

MDA-T32 and MDA-T120 cells were transfected in 96-well microplates and cultured in complete RPMI-1640 medium for 72 h. Confluent cells were starved in L-glutamine- and serum-free DMEM medium (Gibco) for 24 h. Caspase-Glo 3/7 assay (Promega) was used according to the manufacturer’s manual. Measurements were made using Glomax microplate reader (Promega). The final luminescence values were expressed as a ratio between the luminescence of samples and blanks (filled with DMEM). Experiments were repeated three times with technical duplicates.

### Soft agar assay

SeaPlaque agarose (Lonza, Rockland, ME, USA) was mixed with complete RPMI-1640 medium to produce 0.8% base agarose layers in 6-well plates. MDA-T32 and MDA-T120 cells were transfected, cultured for 48 h, re-suspended in complete medium, and mixed with liquid agarose. 1 ml of this suspension (1 × 10^4^ cells in 0.42% agarose) was applied to the agarose-coated wells. The upper agarose layer was covered with 1 ml of complete RPMI-1640 medium (replaced every 5 days). Cells were cultured for 16 (MDA-T32) or 21 days (MDA-T120). Gels were stained with 0.005% crystal violet in 10% ethanol, and photographed. Colonies were counted using ImageJ (NIH, VA, USA). Assays were repeated three times with technical triplicates.

### RNA sequencing

Total RNA was isolated from MDA-T32 cells 72 h after GapmeR transfections (anti-*DIRC3*-common_3, anti-*IGFBP5*, or negative control; all in triplicates). All steps of RNA-seq workflow were performed by GENEWIZ (Leipzig, Germany; see Supplementary Methods and Supp. Table 6). Differential gene expression was determined using DESeq2.^49^ Gene ontology analysis was performed using GeneSCF.^50^

### Stable overexpression of *IGFBP5*

pcDNA3-IGFBP5-V5 plasmid was a gift from Steven Johnson (Addgene plasmid #11608; http://n2t.net/addgene:11608; RRID: Addgene_11608). pcDNA3 plasmid (Invitrogen, Carlsbad, CA, USA) was used as a control vector. Plasmid transfections were performed in MDA-T32 cell line using Fugene 6 (Promega). Transfected cells were cultured for over 3 weeks in complete RPMI-1640 medium containing G418 (600 μg/ml; Clontech, Palo Alto, CA, USA) to select resistant clones.

### IGF-1 stimulation and Western blotting

MDA-T32 cells were transfected with GapmeRs, cultured in complete RPMI-1640 medium for 72 h, and then in serum-free RPMI-1640 containing IGF-1 (20 ng/ml; Gibco) for another 24 h. Next, the conditioned medium was collected and stored at 4 °C, while the cells were serum- and IGF-1-starved for another 12 h. Finally, cells were stimulated for 10 minutes with the conditioned media, each prepared in three versions: a) unmodified, b) enhanced with IGF-1 (extra 30 ng/ml), or c) supplemented with recombinant human IGFBP5 protein (500 ng/ml; Peprotech, Rocky Hill, NJ, USA). Cell lysates were obtained using RIPA buffer with the Halt Protease and Phosphatase Inhibitor Cocktail (Thermo Fisher, Rockford, IL, USA). The conditioned media (1.8 ml) collected in parallel experiments were concentrated using Pierce Protein Concentrator PES, 10K MWCO (Thermo Fisher) and mixed with the RIPA and Inhibitor Cocktail buffer. Electrophoresis was performed using 10% Mini-PROTEAN TGX gel (10 μg of protein per lane). Proteins were transferred to PVDF membranes (both Bio-Rad, Hercules, CA, USA). Membranes were rinsed, blocked with non-fat dry milk, incubated with primary antibodies (Supp. Table 7; 4 °C, overnight), rinsed, and probed with secondary antibodies (room temperature, 1 h). Detection was performed using Clarity Western ECL Substrate (Bio-Rad). Chemidoc Touch System (Bio-Rad) was used for image acquisitions. ImageJ software was used for the densitometric analysis. Experiments were repeated three times.

### Statistical analysis

Statistics were calculated using GraphPad Prism 6 (GraphPad, San Diego, CA, USA). Used tests are shown in Figures. Overlap between DEG sets was analyzed using the hypergeometric distribution calculator (http://nemates.org/MA/progs/overlap_stats.html/).

## List of abbreviations

DTC: differentiated thyroid cancers
cPTC: conventional papillary thyroid cancer
DEG: differentially expressed gene
DIRC3: disrupted in renal carcinoma 3
ERK: extracellular signal-regulated kinase
FBS: fetal bovine serum
FOV: field of view
FTC: follicular thyroid cancer
fvPTC: follicular variant papillary thyroid cancer
GO: gene ontology
GTEx: Genotype-Tissue Expression (project)
GWAS: genome-wide association study
HTC: Hürthle cell thyroid cancer
IGF-1: insulin-like growth factor 1
IGF-1R: insulin-like growth factor 1 receptor
IGFBP5: insulin-like growth factor binding protein 5
lncRNA: long non-coding RNA
PTC: papillary thyroid cancer
SD: standard deviation
SNP: single nucleotide polymorphism
TCGA: The Cancer Genome Atlas (project)

## Acknowledgement

We would like to express our deepest gratitude to Prof. Krystian Jażdżewski for sponsoring the project, providing necessary resources and facilities without which it would not have been possible to complete this study. This work was supported by grants from the National Science Centre, Poland (Preludium grant no. 2017/27/N/NZ2/03116 to P.T.W.), Medical University of Warsaw, Poland (The Young Investigator Grant no. MB/M/12(26) to P.T.W), and Polish Ministry of Education and Science (Regional Initiative for Excellence grant no. 013/RID/2018/19).

## Author Contributions

P.T.W. conceived the project, designed and performed experiments, collected data, performed the analysis, and wrote the manuscript. K.C. performed experiments, collected data, and corrected the manuscript. A.A.M. helped to design experiments, contributed analytic tools, and corrected the manuscript. M.K. participated in the project design, contributed analytic tools, and provided methodological support. D.N. supervised the project, helped to design experiments, contributed analytic tools, and corrected the manuscript.

## Data Availability Statement

The RNA-Seq data generated in this study have been deposited in the Sequence Read Archive (SRA) database with the accession number PRJNA924305. Uncroped images of western blot membranes are shown in the Supplementary Figures 12 and 13. The Cancer Genome Atlas data for Thyroid Carcinoma (PanCancer Atlas) has been obtained using cBioPortal for Cancer Genomics (Memorial Sloan Kettering Cancer Center, USA). User-curated dataset created during the analysis can be accessed here: https://www.cbioportal.org/study/summary?id=thca_tcga_pan_can_atlas_2018#sharedGroups=6187b26ef8f71021ce56ebdd,6187b29bf8f71021ce56ebde. Additional primary data and materials that support the findings of this study are available upon reasonable request.

## Ethics statement

Collection of thyroid tissue was approved by the Institutional Review Board at the Medical University of Warsaw (Aproval KB/184/2009). Written informed consents were obtained from all patients.

