## Supplementary Figures, Tables and Methods for "*Disrupted in Renal Carcinoma 3* (*DIRC3*) impacts malignant phenotype and IGFBP5/IGF-1/Akt signaling axis in differentiated thyroid cancer"

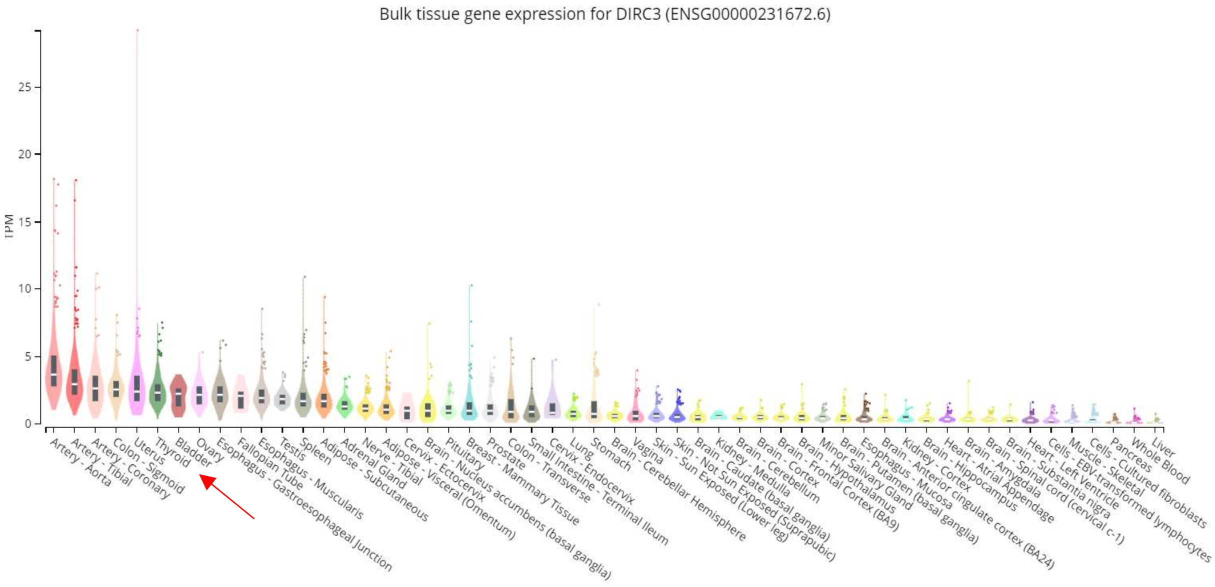

**Supplementary Figure 1.** Expression of *DIRC3* in normal tissues as profiled in the GTEx project (https://gtexportal.org/home/; accessed May 2018). Results for thyroid are marked with a red arrow. Abbreviation: TPM, transcripts per million.

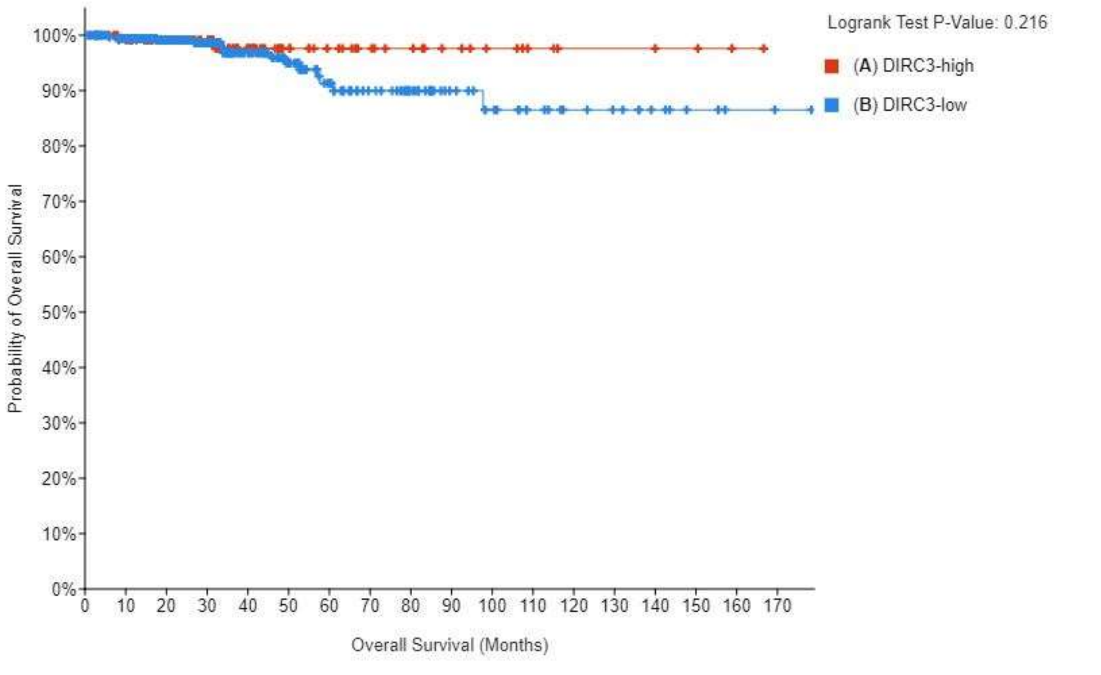

| | Number of Cases, Total | Number of Events | Median Months Overall (95% CI) |
|---|---|---|---|
| (A) DIRC3-high | **126** | **2** | **NA** |
| (B) DIRC3-low | **373** | **14** | **NA** |

**Supplementary Figure 2.** Overall survival of PTC patients according to the *DIRC3* expression status in the TCGA dataset.

**A.**

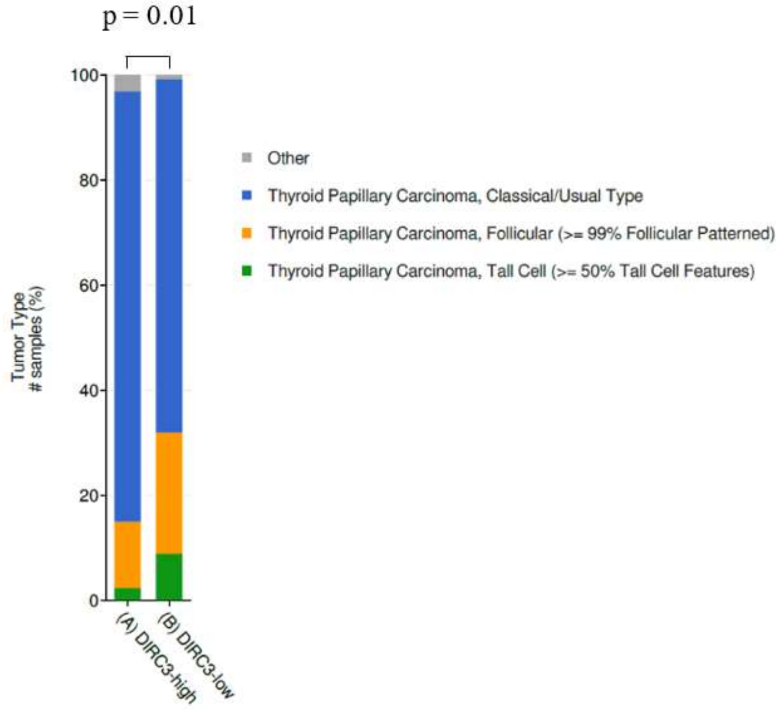

**B.**

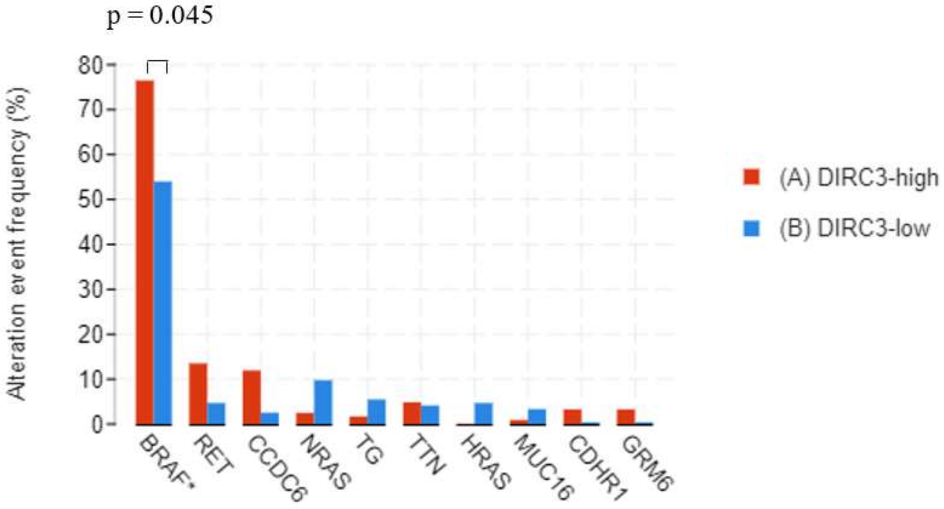

**Supplementary Figure 3.** Histological and molecular characteristics of PTCs in *DIRC3*-high and *DIRC3*-low groups in TCGA data. **(A).** Histological composition of PTCs in *DIRC3*-high and *DIRC3*-low groups (tested with Kruskal-Wallis test with Benjamini-Hochberg correction). **(B).** Frequency of gene mutations in the *DIRC3*-high and *DIRC3*-low PTCs (Fisher Exact test with Benjamini-Hochberg correction). Graphs were prepared using cBioPortal (https://www.cbioportal.org/).

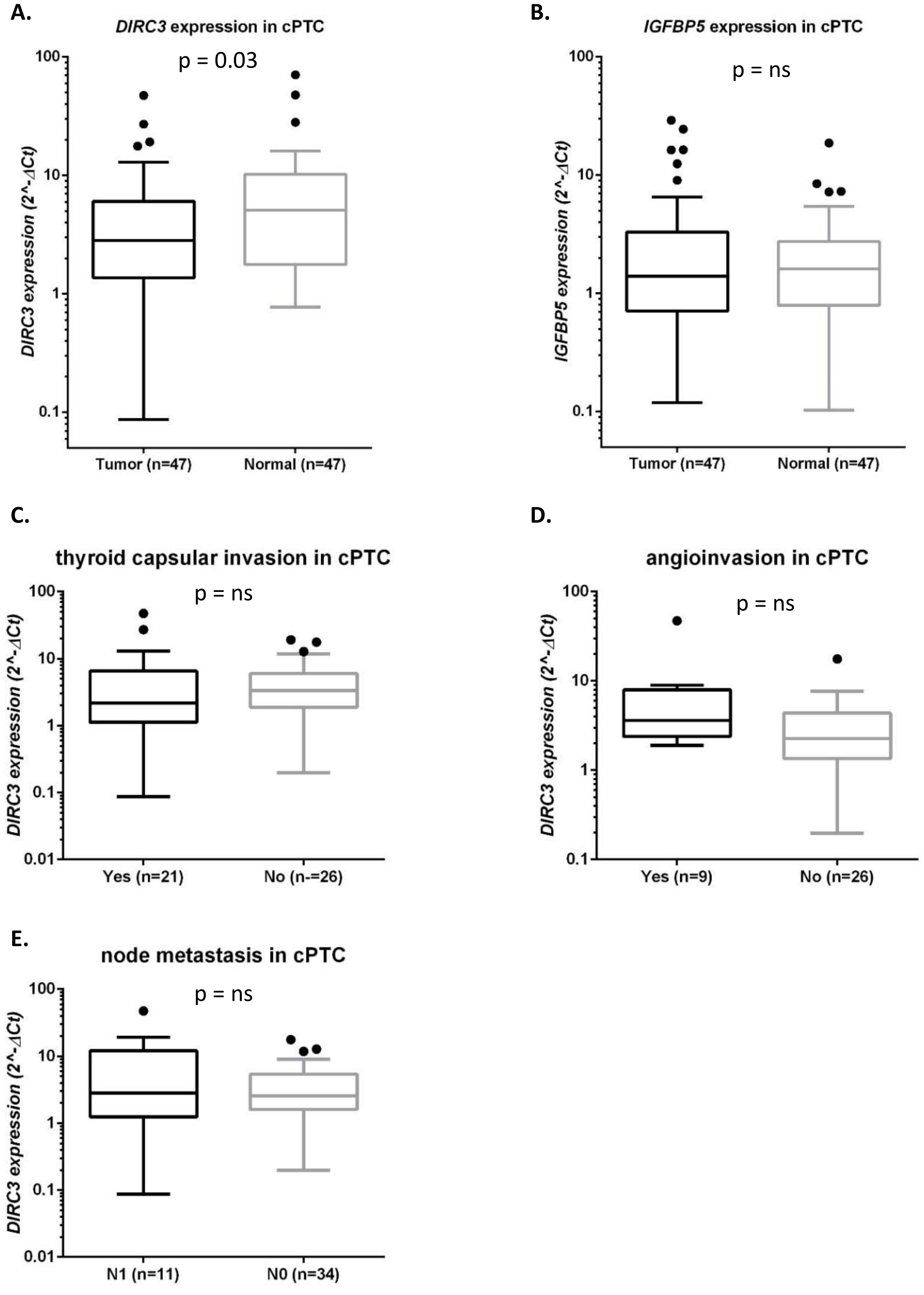

**Supplementary Figure 4.** *DIRC3* expression and clinicopathological features of conventional PTCs. **(A).** Expression of *DIRC3* in conventional PTCs (cPTCs) and the patient-matched normal thyroid tissue. Tested with Wilcoxon test. **(B).** Expression of *IGFBP5* in cPTCs and patient-matched normal thyroid tissue. Tested with Wilcoxon test. **(C).** Expression of *DIRC3* in cPTCs stratified by the presence of capsular invasion, **(D)** angioinvasion, and **(E)** lymph node metastasis. Analyzed with Mann–Whitney U test. Graphs are Tukey plots.

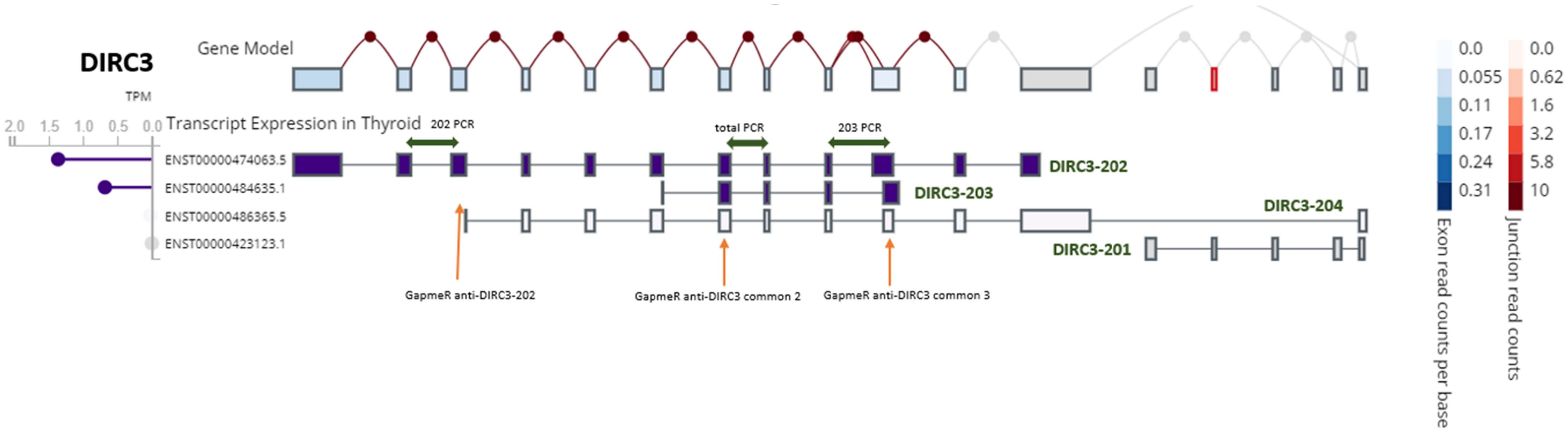

**Supplementary Figure 5.** Structure of *DIRC3* and expression of its splice variants in normal thyroid tissue.

Data was retrieved from the GTEx project via the GTEx Portal (https://gtexportal.org/home/; accessed May 2018).

*DIRC3* is expressed in thyroid in two splice variants: *DIRC3-202* and *DIRC3-203*. qPCR primers were designed to test gene expression (amplicons are shown as green horizontal arrows). Multiple ASOs were designed to silence *DIRC3* (marked as orange vertical arrows).

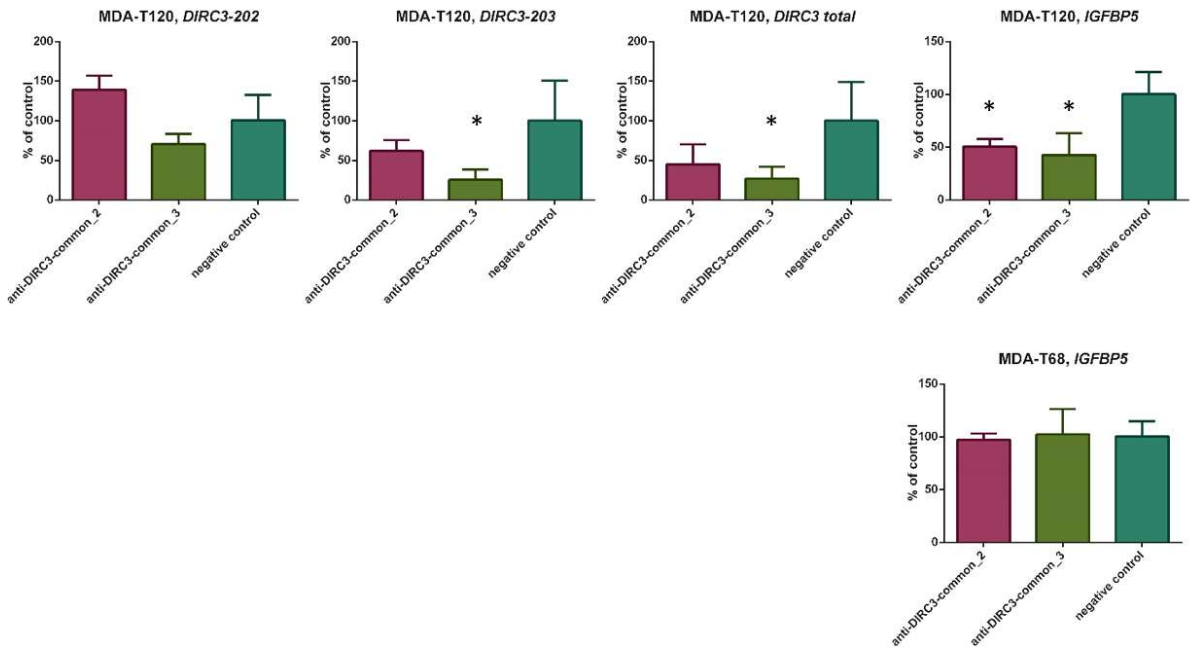

**Supplementary Figure 6**. Silencing of *DIRC3* in MDA-T120 and MDA-T68 cell lines - expression of *DIRC3* total, *DIRC3* splice variants, and *IGFBP5*. Data for MDA-T32 and MCF-7 cell lines is shown in Figure 3C. *DIRC3* was not expressed in MDA-T68 (see Figure 3A). * indicates a significant down-regulation in comparison to the negative control ($p < 0.05$; ANOVA with Dunnett's test). Relative expression values are shown as mean ± SD ($n = 3$).

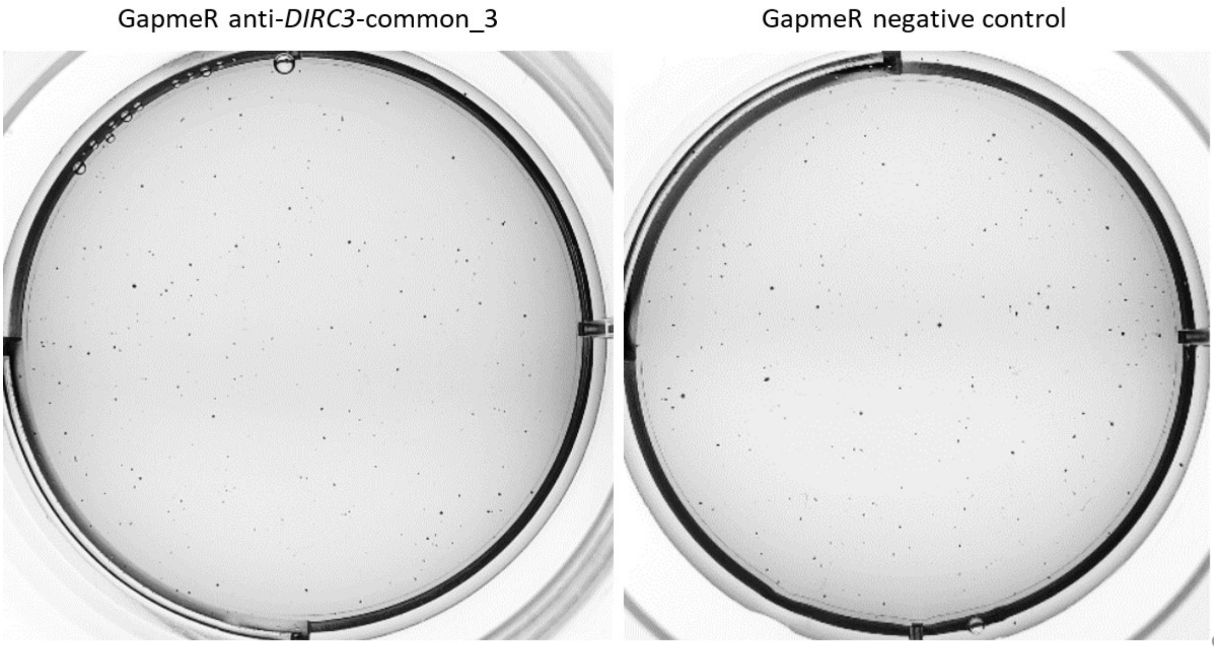

**Supplementary Figure 7.** Anchorage-independent growth of MDA-T32 cells transfected with GapmeRs. Representative wells showing colonies of MDA-T32 cells in soft agar.

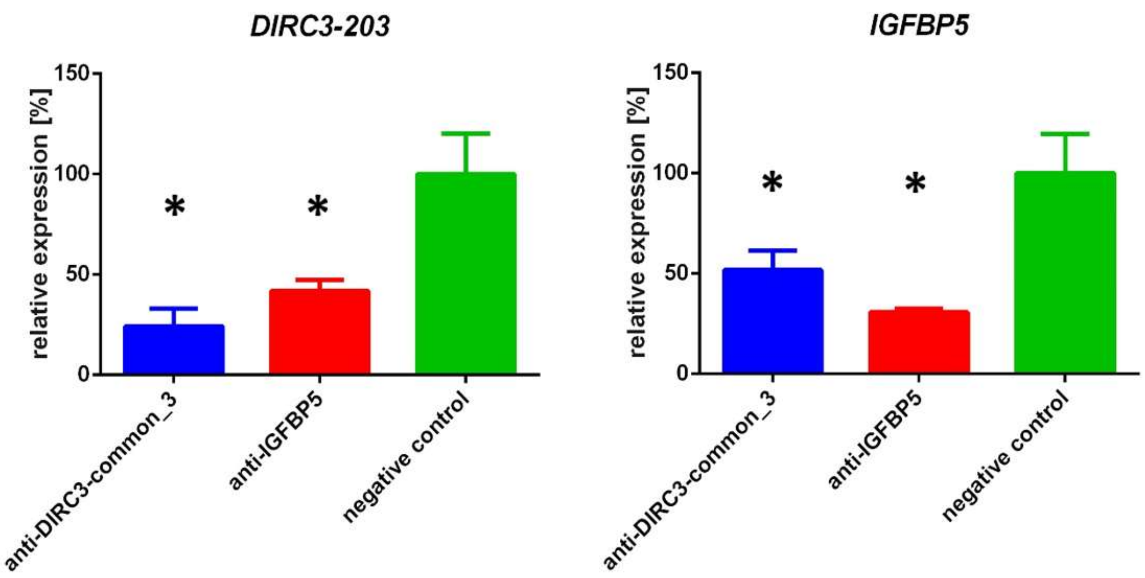

**Supplementary Figure 8.** Silencing of *DIRC3* and *IGFBP5* in MDA-T32 samples prepared for RNA-sequencing. Gene expression was analyzed using qRT-PCR. Transfections were repeated three times (n = 3). Results are shown as mean ± SD. * indicates a significant downregulation as compared to the control samples ($p < 0.05$; ANOVA with Dunnett's test).

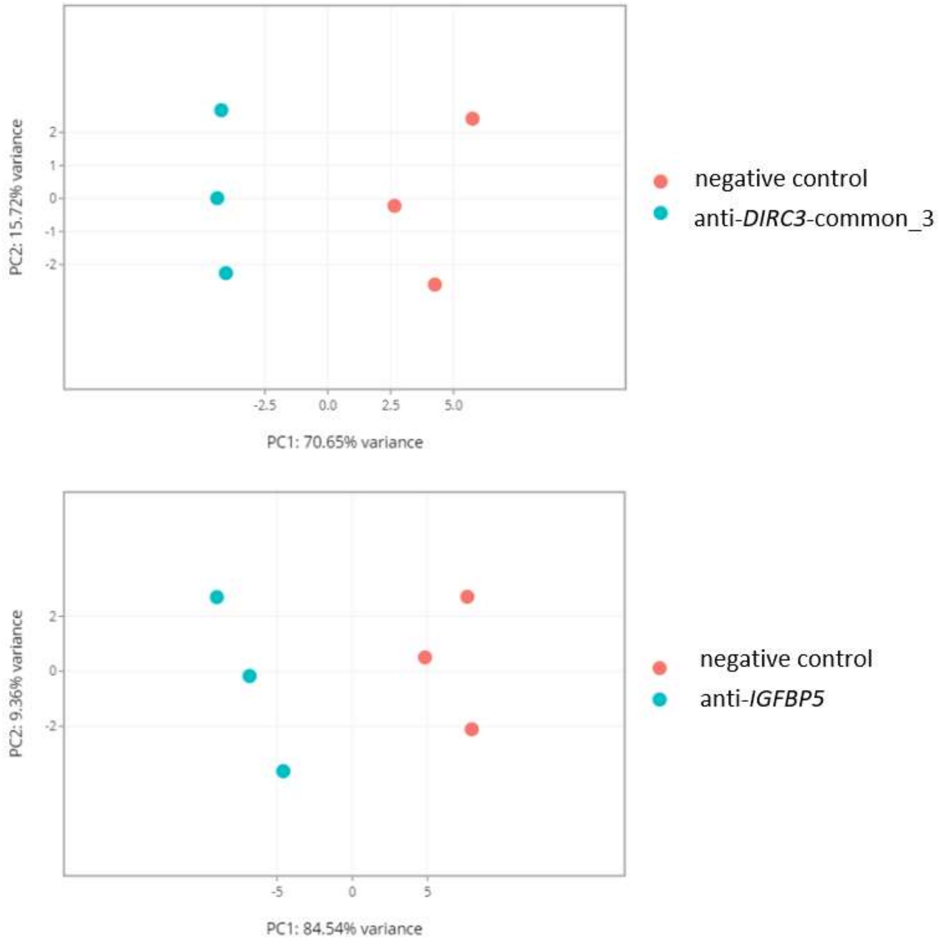

**Supplementary Figure 9.** Principal components analysis (PCA) of the GapmeR-transfected MDA-T32 triplicates in RNA-seq. Cells were transfected with three types of GapmeRs (anti-*DIRC3*-common_3, anti-*IGFBP5*, or negative control).

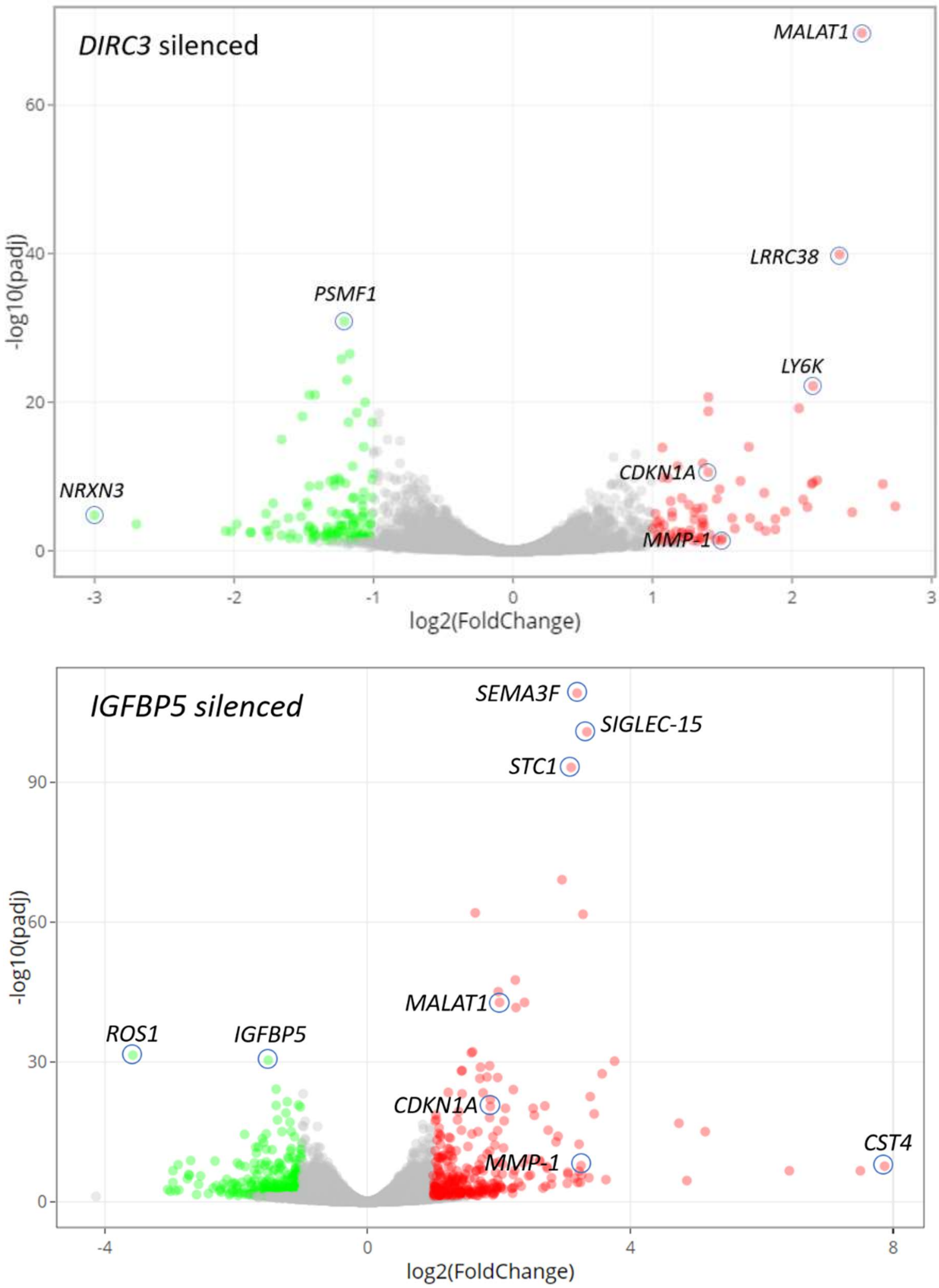

**Supplementary Figure 10.** Volcano plots illustrating changes in the gene expression profile after silencing of *DIRC3* or *IGFBP5* in MDA-T32 cells. DEGs are shown as color dots (red: upregulated, green: downregulated). Certain prominent DEGs are marked.

A.

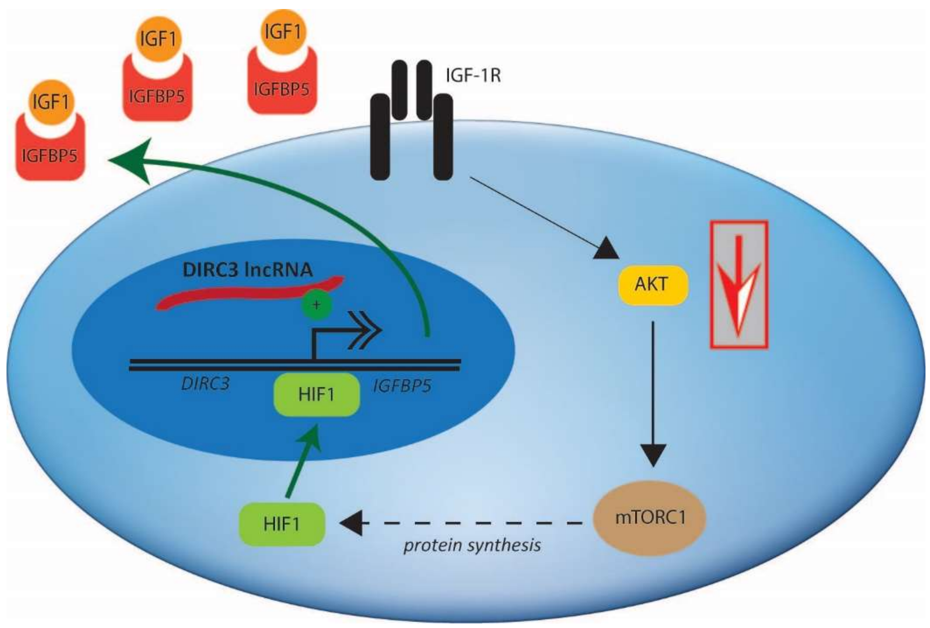

B.

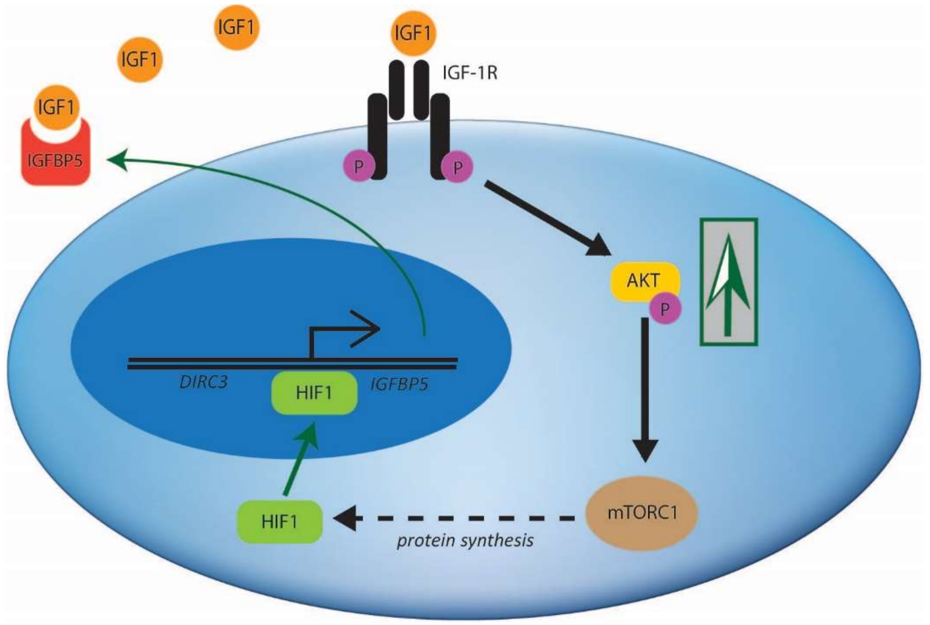

**Supplementary Figure 11.** Model of the putative role of *DIRC3* in the IGF-1 signaling pathway in thyroid cancer cells. The model is based on literature data and the results of this study. **(A).** IGF-1R is stimulated by IGF-1, what promotes signaling cascade involving Akt and mammalian target of rapamycin complex 1 (mTORC1). Augmented flux in the Akt signaling pathway drives cancer progression. Additionally, mTORC1 upregulates protein synthesis. Among the proteins upregulated by IGF-1 are components of hypoxia-inducible factor 1 (HIF-1), a transcription factor promoting expression of many oncogenes.[1] One of transcriptomic targets of HIF-1 is *IGFBP5*, which is consequently upregulated.[2-5] In thyroid cells that retain adequate expression of *DIRC3*, this lncRNA further promotes expression of *IGFBP5*. Extracellular release of IGFBP5 limits the bioavailability of IGF-1 and prevents its excessive stimulatory effect.[1, 6] Accordingly, a negative feedback loop is formed. **(B).** Production of IGFBP5 is downregulated in thyroid cancers with low expression of *DIRC3*. Accordingly, the bioavailability of IGF-1 is higher and the oncogenic Akt signaling is augmented. The negative feedback loop is also impaired. Arrow thickness in the schemes indicates the magnitude of signaling flow.

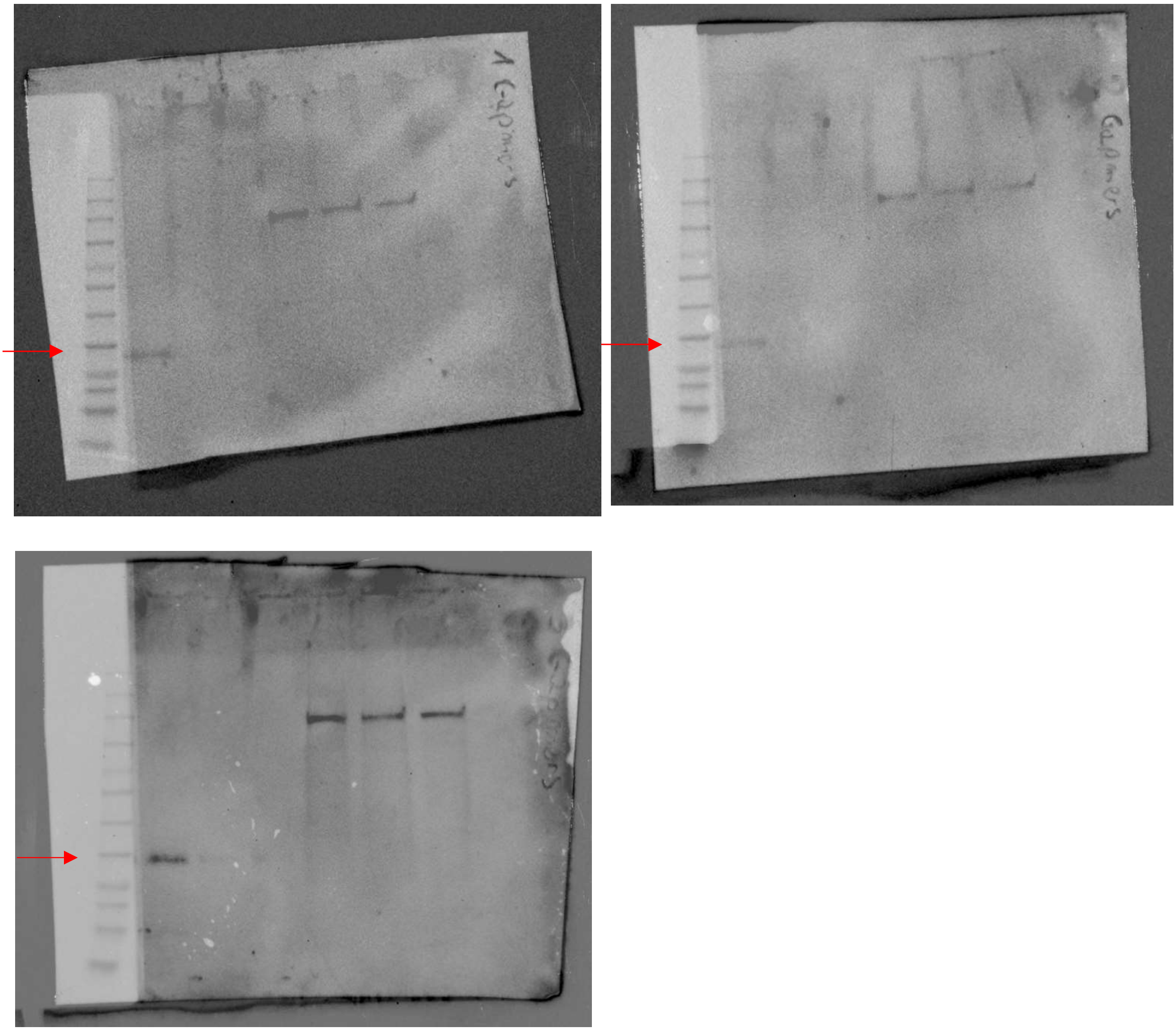

**Supplementary Figure 12.** Immunoblot membranes demonstrating detection of IGFPB5 (32 kDa; red arrow) in the GapmeR-transfected MDA-T32 cells cultured for 24 h with IGF-1 (three biological replicates). The sample order: 1. conditioned medium, negative control; 2. conditioned medium, *IGFBP5* silenced; 3. conditioned medium, *DIRC3* silenced; 4. cell lysate, negative control; 5. cell lysate, *IGFBP5* silenced; 6. conditioned medium, *DIRC3* silenced. The ladder: Perfect Tricolor Protein Ladder (EURx, Gdansk, Poland).

**A.**

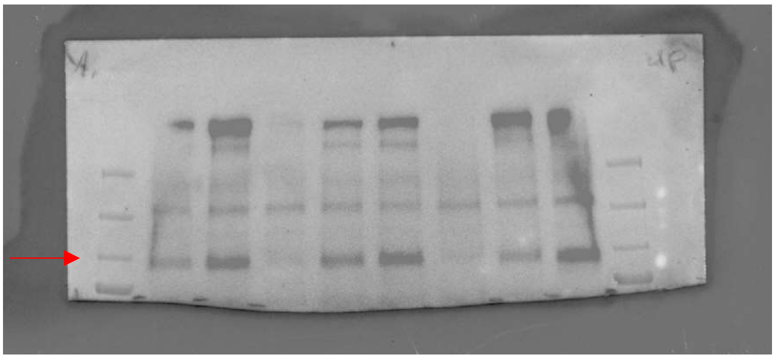

**B.**

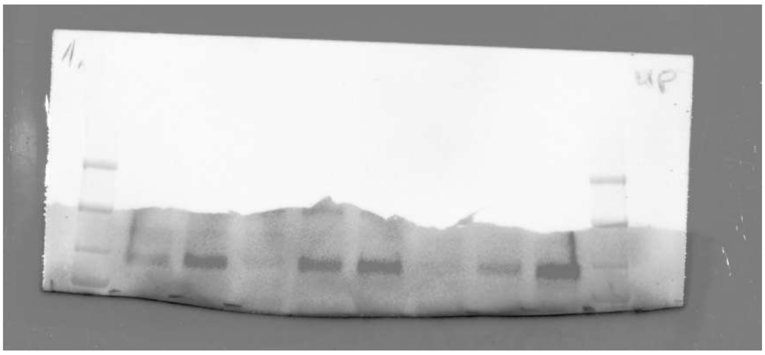

**C.**

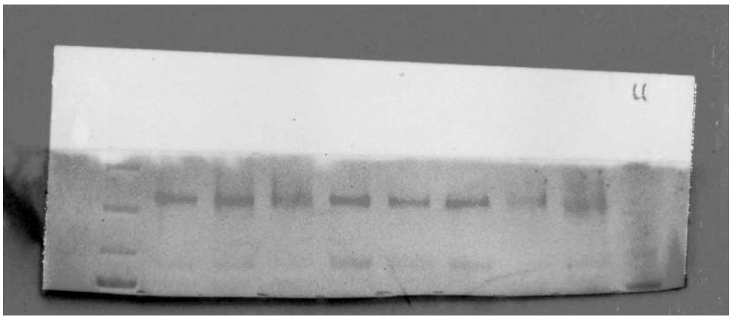

**D.**

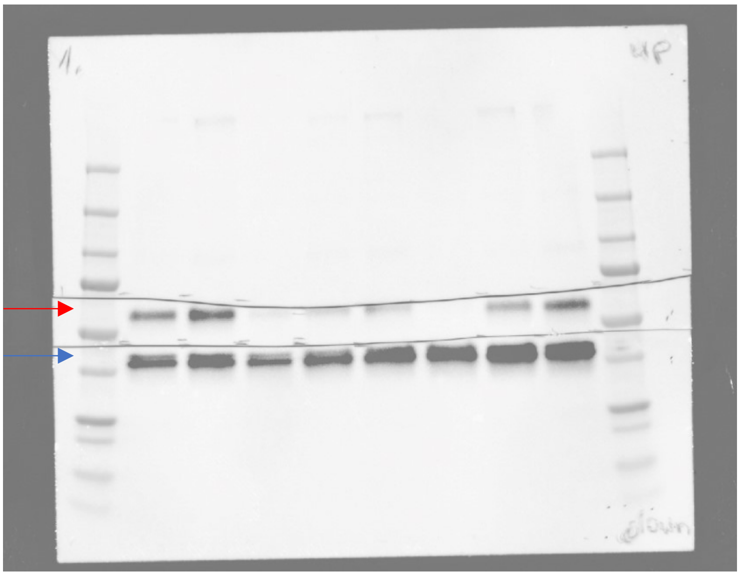

**E.**

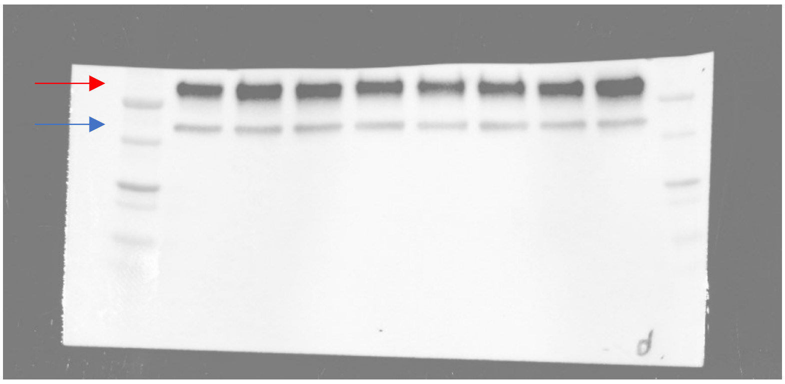

**F.**

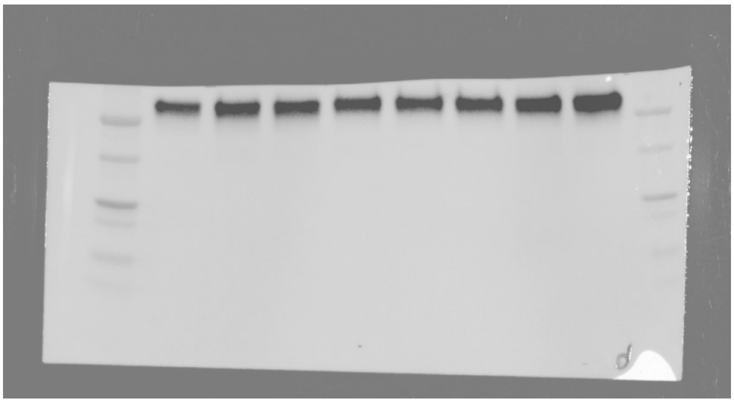

**G.**

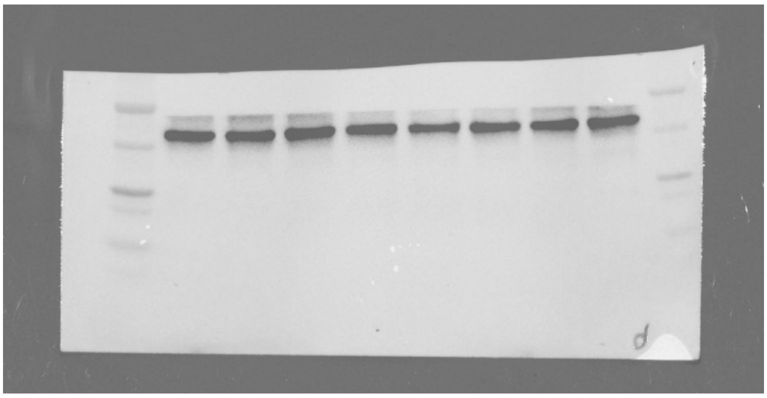

**Supplementary Figure 13.** Immunoblot membranes demonstrating detection of proteins in the GapmeR-transfected MDA-T32 cells stimulated with conditioned media (replicate 1). The sample order in all membranes: 1. *DIRC3* silenced, conditioned medium (unmodified, i.e. with 20 ng/ml of IGF-1); 2. *DIRC3* silenced, conditioned medium with extra 30 ng/ml of IGF-1; 3. *DIRC3* silenced, conditioned medium with recombinant human IGFBP5 (500 ng/ml); 4. Negative control GapmeR, conditioned medium (unmodified, i.e. with 20 ng/ml of IGF-1); 5. Negative control GapmeR, conditioned medium with extra 30 ng/ml of IGF-1; 6. Negative control GapmeR, conditioned medium with recombinant human IGFBP5 (500 ng/ml); 7. *IGFBP5* silenced, conditioned medium (unmodified, i.e. with 20 ng/ml of IGF-1); 8. *IGFBP5* silenced, conditioned medium with extra 30 ng/ml of IGF-1. The ladder: Precision Plus Protein Dual Color Standards (Bio-Rad, Hercules, CA, USA). **(A).** pIGF-1R β (95 kDa). **(B).** pIGF-1R β (95 kDa), different exposure **(C).** IGF-I Receptor β (95 kDa); **(D).** pAKT (60 kDa, red arrow) and pERK1/2 (44, 42 kDa, blue arrow). **(E).** Akt (pan) (60 kDa, red arrow) and β-actin (45 kDa, blue arrow). **(F).** Akt (pan) (60 kDa), different exposure. **(G).** β-actin (45 kDa), different exposure.

### SUPPLEMENTARY TABLES

| Gene | Cytoband | Spearman's correlation | p-value |
|---|---|---|---|
| ***IGFBP5*** | 2q35 | 0.79 | 1.61E-103 |
| ***DMD*** | Xp21.2-p21.1 | 0.64 | 2.21E-56 |
| ***LCN6*** | 9q34.3 | 0.59 | 2.82E-46 |
| ***QSOX2*** | 9q34.3 | 0.57 | 2.70E-43 |
| ***FIGNL2*** | 12q13.13 | 0.56 | 3.86E-41 |
| ***ESRRA*** | 11q13.1 | -0.56 | 5.49E-41 |
| ***WSB2*** | 12q24.23 | -0.56 | 1.31E-40 |
| ***CDK5RAP2*** | 9q33.2 | 0.55 | 4.31E-40 |
| ***ADTRP*** | 6p24.1 | 0.55 | 1.04E-39 |
| ***SCNN1G*** | 16p12.2 | 0.55 | 3.39E-39 |
| ***PTK7*** | 6p21.1 | 0.55 | 6.37E-39 |
| ***PPM1H*** | 12q14.1-q14.2 | -0.55 | 1.41E-38 |
| ***USP27X*** | Xp11.23 | 0.54 | 2.04E-38 |
| ***ZFTA*** | 11q13.1 | 0.54 | 4.53E-38 |
| ***INMT*** | 7p14.3 | 0.54 | 5.91E-38 |
| ***LCN8*** | 9q34.3 | 0.54 | 6.56E-38 |
| ***PLAAT1*** | 3q29 | -0.54 | 8.20E-38 |

**Supplementary Table 1.** List of genes demonstrating the strongest expression correlations with *DIRC3* in PTCs (TCGA data).

| | |
|---|---|
| **Sex** | Female, n = 55<br>Male, n = 10<br>data missing, n = 2 |
| **Age at the time of diagnosis** | mean 48.1 years (standard deviation: 14.5 y);<br>range 18-80 years |
| **Histological type** | PTC, conventional variant, n = 47<br>PTC, follicular variant, n = 9<br>FTC, n = 5<br>FTC + follicular variant of PTC, n = 1<br>Hürthle cell cancer, n = 3<br>data missing (DTC), n = 2 |
| **Multifocality** | Yes, n = 15<br>No, n = 50<br>data missing, n = 2 |
| **Angioinvasion** | Yes, n = 14<br>No, n = 33<br>data missing, n = 20 |
| **Invasion of thyroid capsule** | Yes, n = 23<br>No, n = 38<br>data missing, n = 7 |
| **T stage<br>(AJCC/TNM 7th Edition)** | pT1a, n = 25<br>pT1b, n = 19<br>pT2, n = 6<br>pT3, n = 15<br>data missing, n = 2 |
| **N stage<br>(nodal metastases)** | N0, n = 44<br>N1, n = 13<br>data missing, n = 10 |
| **M stage<br>(distant metastases)** | M0, n = 53<br>M1, n = 4<br>data missing, n = 10 |

**Supplementary Table 2.** Clinical and pathological information on DTC patients included in the study cohort.

| GENE | FULL NAME | EVIDENCE FOR ROLE IN THYROID CANCERS | POSSIBLE ROLE | EXPRESSION UPON *DIRC3* SILENCING |
|---|---|---|---|---|
| ***NRXN3*** | *Neurexin 3-alpha* | | | ↓ |
| ***COL17A1*** | *Collagen Type XVII Alpha 1 Chain* | Downregulated in PTCs and anaplastic thyroid cancers as compared to normal thyroid tissue.[1, 2] | | ↑ |
| ***WNT11*** | *Wnt Family Member 11* | | | ↓ |
| ***FER1L4*** | *Fer-1 Like Family Member 4, Pseudogene (Functional)* | Upregulation of *FER1L4* correlated positively with the lymph node metastasis, extrathyroidal extension and advanced TNM stages in PTCs. Knockdown of *FER1L4* suppressed cell proliferation, migration and invasion in PTCs, whereas ectopic expression of *FER1L4* promoted these processes.[3] | Oncogene | ↓ |
| ***ANGPT2*** | *Angiopoietin-2* | Expression of *ANGPT2* gradually increased from non-neoplastic thyroid tissue to high-risk thyroid cancers.<br>*ANGPT2* expression was useful to distinguish cancers from benign thyroid neoplasms.<br>Expression of *ANGPT2* was predictive of the lenvatinib sensitivity in radioiodine-refractory PTCs.[4-7] | Possible oncogene | ↑ |
| ***DBP*** | *D-box binding PAR bZIP transcription factor* | | | ↓ |
| ***TNFSF8*** | *Tumor Necrosis Factor (Ligand) Superfamily, Member 8, CD30L* | The proportion of CD30L+ cells correlated inversely with the PTC aggressiveness.[8] | | ↑ |
| ***AREG*** | *Amphiregulin* | Overexpressed in PTCs.<br>Upregulation of *AREG* associated with worse prognosis and increased risk of cervical node metastasis in PTCs.[9-11] | Possible oncogene | ↑ |
| ***MGP*** | *Matrix Gla Protein* | | | ↓ |
| ***IL1RL1*** | *Interleukin 1 Receptor Like 1* | | | ↑ |
| ***ACTL8*** | *Actin Like 8* | | | ↑ |
| ***HSD11B1*** | *Hydroxysteroid 11-Beta Dehydrogenase 1* | | | ↑ |

| | | | | |
|---|---|---|---|---|
| ***CDKN1A*** | *Cyclin Dependent Kinase Inhibitor 1A, p21* | Inhibits proliferation and arrests the cell cycle.<br>*CDKN1A* deletion is involved in thyroid carcinogenesis.<br>The expression level of *CDKN1A* was useful to distinguish adenomas from PTCs.<br>Progressive loss of p21 expression was observed with advancing clinical stage of DTCs. Expression of p21 was higher in large PTCs and in tumors extending beyond the thyroid capsule.<br>No correlation between the p21 expression, clinical parameters and patients' prognosis could be established in one study.[12-20] | Possibly tumor suppressor | ↑ |
| ***BEX1*** | *Brain Expressed X-Linked 1* | Downregulated in thyroid cancers.[21, 22] | | ↑ |
| ***FST*** | *Follistatin* | FST serum level was increased in thyroid cancer patients. Its level correlated positively with the cancer aggressiveness (metastasis, vascular invasion, TNM).[23] | | ↑ |
| ***DGKB*** | *Diacylglycerol Kinase Beta* | | | ↓ |
| ***CYP1B1*** | *Cytochrome P450 Family 1 Subfamily B Member 1* | Expression of *CYP1B1* was elevated in PTCs, but was lower in anaplastic cancer as compared to normal thyroid tissue.[22, 24, 25] | | ↓ |
| ***FBLN7*** | *Fibulin 7* | Downregulated in PTCs as compared to normal thyroid tissue.[25-27] | | ↓ |
| ***ST8SIA6*** | *Alpha-2,8-sialyltransferase* | Downregulated in FTCs. *ST8SIA6* inhibited cell proliferation, migration and invasion in FTCs, while its upregulation decreased the cancer invasiveness.[28] | Tumor suppressor | ↓ |
| ***FAM13C*** | *Family With Sequence Similarity 13 Member C* | | | ↓ |
| ***LRMDA*** | *Leucine Rich Melanocyte Differentiation Associated* | | | ↓ |
| ***ROBO4*** | *Roundabout guidance receptor 4* | Overexpressed in PTCs.<br>Iodine therapy might exert therapeutic effects by targeting *ROBO4* in thyroid cancer cells.[29, 30] | | ↑ |
| ***PPP2R2B*** | *Protein Phosphatase 2 Regulatory Subunit beta* | Downregulated in malignant thyroid nodules.[31, 32] | | ↓ |
| ***STC1*** | *Stanniocalcin 1* | Silencing of *STC1* inhibited the thyroid cancer cell proliferation, whereas the recombinant human STC1 enhanced cell proliferation.<br>Altered expression in thyroid tumor cell lines and tumor tissues.<br>Overexpression was a negative prognostic biomarker in the thyroid cancer patients.[33-37] | Oncogene | ↑ |

| | | | | |
|---|---|---|---|---|
| ***LY6K*** | *Lymphocyte Antigen 6 Family Member K* | | | ↑ |
| ***LRRC38*** | *Leucine Rich Repeat Containing 38* | | | ↑ |
| ***IL24*** | *Interleukin 24* | Involved in the cell cycle regulation and immune suppression. Expression induced by *RET/PTC* oncogene in PTCs. Induction of IL24 by oncogenes may support cell proliferation and tumor growth at the early stages of thyroid carcinogenesis.[38, 39] | Possibly oncogene | ↑ |
| ***FAM198B*** | *Golgi Associated Kinase 1B* | | | ↓ |
| ***ESM1*** | *Endothelial Cell Specific Molecule 1* | Expression elevated in the radiation-induced PTCs as compared to sporadic PTCs.[40] Expression of *ESM1* was increased in FTCs harboring *PAX8-PPARγ1* rearrangement. *ESM1* may impact cell adhesion and angiogenesis.[41] *ESM1* was differentially expressed in papillary thyroid microcarcinoma and PTCs.[42] | | ↑ |
| ***MYO5B*** | *Myosin VB* | Expression of *MYO5B* was induced by the thyroid transcription factor PAX8. *MYO5B* may regulate the follicular polarity in thyroid cells[43].*MYO5B* was downregulated in anaplastic thyroid cancers.[44] | | ↑ |
| ***ACTBL2*** | *Actin Beta Like 2* | Expression of *ACTBL2* was lower in the follicular variant PTCs than in convention PTCs.[45] | | ↓ |
| ***ADRB2*** | *Adrenoceptor Beta 2* | Expression of *ADRB2* was higher in DTCs than in normal thyroid tissue.[46] | | ↑ |
| ***SSC5D*** | *Scavenger Receptor Cysteine Rich Family Member With 5 Domains* | | | ↓ |
| ***TNFSF15*** | *TNF Superfamily Member 15; Vascular Endothelial Cell Growth Inhibitor* | Downregulated by *EPB41L4A* (a validated PTC tumor suppressor). *TNFSF15* was involved in the proliferation and differentiation of thyroid cancer cells.[47] | | ↑ |
| ***BGN*** | *Biglycan* | Expression of *BGN* was higher in PTCs than in oncocytic carcinomas.[48] | | ↓ |
| ***EFHC2*** | *EF-Hand Domain Containing 2* | | | ↓ |
| ***OLFML2A*** | *Olfactomedin Like 2A* | Expression of *OLFML2A* was lower in FTCs than in thyroid adenomas.[49] | | ↓ |

| | | | | |
|---|---|---|---|---|
| ***WDR86*** | *WD Repeat Domain 86* | | | ↓ |
| ***SAMD11*** | *Sterile alpha motif domain-containing protein 11* | | | ↓ |
| ***INSC*** | *Inscuteable Spindle Orientation Adaptor Protein* | | | ↑ |
| ***VSTM1*** | *V-Set And Transmembrane Domain Containing 1* | *VSTM1* expression impacts the radiation sensitivity of anaplastic and poorly differentiated thyroid cancers.[50] | | ↑ |
| ***MMP1*** | *Matrix Metallopeptidase 1* | High *MMP-1* expression was observed in undifferentiated thyroid cancer cell lines.<br>MMP-1 level was significantly higher in follicular thyroid carcinomas than in adenomas. High expression of MMP-1 in DTCs correlated positively with the cancer aggressiveness (laryngotracheal invasion, multifocality, regional and distant metastases).<br>Elevated expression of MMP-1 in DTCs was a risk factor for cervical metastases.<br>No relationship between the MMP-1 expression in DTCs and the cancer invasiveness, metastasis, or recurrence was detected in another study[51-58]. | Oncogene | ↑ |
| ***SERPINB2*** | *Serpin Family B Member 2; Plasminogen Activator Inhibitor 2* | *SERPINB2* inhibited apoptosis in anaplastic thyroid cancer.[59] | | ↑ |
| ***TRIM15*** | *Tripartite motif-containing 15* | | | ↓ |
| ***AL162231.2*** | - | | | ↓ |
| ***AL512488.1*** | | | | ↑ |
| ***AC121764.1*** | - | | | ↓ |
| ***AC091173.1*** | - | | | ↑ |
| ***MALAT1*** | *Metastasis Associated Lung Adenocarcinoma Transcript 1* | *MALAT1* promoted cell proliferation, migration, invasiveness and angiogenesis in thyroid cancers.<br>Knockdown of *MALAT1* reduced cell migration and invasiveness in B-CPAP and K1 thyroid cancer cell lines.<br>*MALAT1* upregulated *IGF2BP2* and enhanced expression of *MYC*, conferring a stimulatory effect on the proliferation, migration and invasiveness of thyroid cancer cells. | Oncogene | ↑ |

| | | | | |
|---|---|---|---|---|
| | | *MALAT1* was highly expressed in normal thyroid tissues and in thyroid tumors. Expression of *MALAT1* was higher in PTCs and Hürthle cell thyroid neoplasms than in normal thyroid tissue. Its expression correlated positively with the tumor size, lymph node metastases and disease stage in PTCs.<br>*MALAT1* was, however, downregulated in poorly differentiated thyroid cancers and anaplastic thyroid carcinomas.[60-67] | | |
| ***TRIL*** | *TLR4 Interactor With Leucine Rich Repeats* | | | ↓ |
| ***LINC02407*** | - | | | ↑ |
| ***LINC00052*** | - | | | ↓ |
| ***AC107398.3*** | - | | | ↓ |
| ***AP000894.2*** | - | | | ↓ |
| ***AC004585.1*** | *Immune related lncRNA* | | | ↑ |
| ***DACH1*** | *Dachshund Family Transcription Factor 1* | Indirect evidence for an anti-proliferative role of *DACH1* in PTCs.[68] | | ↓ |
| ***FO681492.1*** | - | | | ↓ |
| ***AC002384.1*** | - | | | ↓ |
| ***AL662907.1*** | | | | ↓ |
| ***BCL2A1*** | *BCL2 Related Protein A* | *BCL2A1* increased the apoptosis resistance in poorly differentiated thyroid cancers.[69] | | ↑ |
| ***KNOP1P5*** | *Lysine Rich Nucleolar Protein 1 Pseudogene 5* | | | ↓ |
| ***STARD4-AS1*** | *STARD4 Antisense RNA 1* | | | ↓ |
| ***CYP1B1-AS1*** | *CYP1B1 antisense RNA 1* | Upregulated in PTCs. Expression of *CYP1B1-AS1* may have a prognostic value.[70, 71] | | ↓ |
| ***MAF*** | *MAF BZIP Transcription Factor; V-Maf Avian Musculoaponeurotic Fibrosarcoma Oncogene* | | | ↑ |
| ***SLC26A9*** | *Solute Carrier Family 26 Member 9* | | | ↓ |

| | | | | |
|---|---|---|---|---|
| ***MYO7B*** | *Myosin VIIB* | | | ↑ |
| ***NPR1*** | *Natriuretic Peptide Receptor 1* | Low expression of *NPR1* was associated with increased risk of PTC recurrence.[72] | | ↓ |
| ***CSF2*** | *Colony Stimulating Factor 2 (GM-CSF)* | Upregulated in PTCs. Expression of *CSF2* was induced by the *RET/PTC3* oncogene.[73-78] | | ↑ |
| ***MEGF6*** | *Multiple epidermal growth factor-like domains protein 6* | | | ↓ |
| ***PART1*** | *Prostate Androgen-Regulated Transcript 1* | | | ↓ |
| ***LRRC4C*** | *Leucine Rich Repeat Containing 4C* | Upregulated in widely-invasive Hürthle cell carcinomas as compared to normal thyroid tissue.[79] | | ↑ |
| ***WDR31*** | *WD Repeat Domain 31* | | | ↓ |
| ***FAM83A*** | *Family With Sequence Similarity 83 Member A* | | | ↑ |
| ***N4BP3*** | *NEDD4 Binding Protein 3* | Downregulated in widely-invasive Hürthle cell carcinomas as compared to normal thyroid tissue.[79] | | ↑ |
| ***ALPK3*** | *Alpha Kinase 3* | | | ↓ |
| ***PTGER2*** | *Prostaglandin E Receptor 2* | | | ↓ |
| ***RIMS3*** | *Regulating Synaptic Membrane Exocytosis 3* | | | ↓ |
| ***CCL26*** | *C-C Motif Chemokine Ligand 26* | | | ↑ |

**Supplementary Table 3.** List of DEGs shared by *DIRC3* and *IGFBP5* in RNA-seq in the gene silencing experiments in MDA-T32 cell line. Literature evidence on the role of particular genes in thyroid carcinogenesis is provided.

| | **Forward primer** | **Reverse primer** |
|---|---|---|
| ***DIRC3*, total (qPCR)** | CTGGTGTCACATCAGAATCTG | ACAATCAGCATGGGACTCAC |
| ***DIRC3-202* (qPCR)** | GGAATTGTGCGAGTGGAAATG | TGCTAAGATAGAGCCGCAAAG |
| ***DIRC3-203* (qPCR)** | TCATGCTGGCCGATGCTGAG | CCACTGGGACAACCATCTTTTAATCAG |
| ***DIRC3-204* (qPCR)** | CTCATCTGTCCGACGAAGCA | CCCTACTGTCCTGGTGGAGA |
| ***IGFBP5* (qPCR)** | AAGATCGAGAGAGACTCCCGT | CCGACAAACTTGGACTGGGT |
| ***HPRT1* (qPCR)** | CAGAGGGCTACAATGTGATG | TGGCGTCGTGATTAGTGATG |
| ***MALAT1* (qPCR)** | GACGGAGGTTGAGATGAAGC | ATTCGGGGCTCTGTAGTCCT |
| ***SNHG5* (qPCR)** | GTGGACGAGTAGCCAGTGAA | GCCTCTATCAATGGGCAGACA |

**Supplementary Table 4.** Primers used in the SYBR Green qPCR assays.

| **GapmeR** | **Sequence** |
|---|---|
| **anti-*DIRC3*-common_2** | T*A*A*A*A*T*G*G*C*A*G*G*G*T*G*T |
| **anti-*DIRC3*-common_3** | C*T*T*G*A*A*T*A*G*A*G*G*C*T*A |
| **anti-*DIRC3*-202** | C*A*C*A*A*A*T*A*A*G*T*A*G*A*C*G |
| **anti-*IGFBP5*** | G*G*A*A*T*G*G*T*G*C*G*A*G*T*A*T |
| **anti-*MALAT1*** | C*G*T*T*A*A*C*T*A*G*G*C*T*T*T*A |
| **negative control A** | A*A*C*A*C*G*T*C*T*A*T*A*C*G*C |

**Supplementary Table 5.** GapmeR oligonucleotides used in the study. * indicates a phosphorothioate backbone modification.

| Sample ID | Total number of sequencing reads | Total number of uniquely mapped reads | RNA quality number (RQN) | Radio of exon-mapped reads to total uniquely mapped reads (Expression Profile Efficiency) | Total number of detected transcripts with reads ≥1 |
|---|---|---|---|---|---|
| T32-Gap-DIRC3-rep5 | 18,374,312 | 18,102,323 | 10 | 0.875 | 23,744 |
| T32-Gap-IGFBP5-rep5 | 20,353,969 | 20,072,937 | 10 | 0.859 | 23,966 |
| T32-Gap-neg-ctr-rep5 | 33,010,474 | 31,668,710 | 10 | 0.881 | 26,044 |
| T32-Gap-DIRC3-rep6 | 22,663,247 | 22,345,632 | 10 | 0.843 | 24,396 |
| T32-Gap-IGFBP5-rep6 | 21,588,483 | 21,312,992 | 10 | 0.836 | 24,154 |
| T32-Gap-neg-ctr-rep6 | 25,542,132 | 25,165,77 | 10 | 0.838 | 25,214 |
| T32-Gap-DIRC3-rep7 | 19,786,136 | 19,522,687 | 10 | 0.822 | 23,700 |
| T32-Gap-IGFBP5-rep7 | 20,127,385 | 19,857,792 | 10 | 0.799 | 23,488 |
| T32-Gap-neg-ctr-rep7 | 27,033,448 | 26,647,369 | 10 | 0.822 | 25,647 |

**Supplementary Table 6.** Detailed information on the quality of mapping of RNA-seq sample reads. Reads were mapped to the Homo sapiens GRCh38 reference genome.

| Antibody | Dilution | Catalog number | Manufacturer |
|---|---|---|---|
| **IGFBP5 (E3H3Z) Rabbit mAb; primary** | 1:1000 | #76541 | Cell Signaling, USA |
| **Phospho-IGF-I Receptor β (Tyr1135) (DA7A8) Rabbit mAb; primary** | 1:1000 | #3918 | Cell Signaling, USA |
| **IGF-I Receptor β (D23H3) XP Rabbit mAb; primary** | 1:1000 | #9750 | Cell Signaling, USA |
| **Phospho-AKT (Ser473) (D9E) XP Rabbit; primary** | 1:2000 | #4060 | Cell Signaling, USA |
| **AKT (pan) (C67E7) Rabbit mAb; primary** | 1:1000 | #4691 | Cell Signaling, USA |
| **Phospho-p44/42 MAPK (Erk1/2) (Thr202/Tyr204) (D13.14.4E) XP Rabbit mAb; primary** | 1:1000 | #4370 | Cell Signaling, USA |
| **β-Actin Antibody, Rabbit mAb; primary** | 1:1000 | #4967 | Cell Signaling, USA |
| **Peroxidase AffiniPure Goat Anti-Rabbit IgG [H+L]; secondary** | 1:10,000 | 111-035-144 | Jackson Immuno Research Labs, USA |

**Supplementary Table 7.** Antibodies used for Western blotting.

### SUPPLEMENTARY METHODS

#### RNA sequencing (library preparation, sequencing and data analysis).

All steps of Next Generation Sequencing (including the RNA quality control, library preparati on,sequencing, bioinformatic analysis) were performed by GENEWIZ (Leipzig, Germany) usingthe "Standard RNA-seq service".

RNA Library Preparation and NovaSeq Sequencing RNA samples were quantified using Qubit 4.0 Fluorometer (Life Technologies, Carlsbad, CA, USA) and RNA integrity was checked with RNA Kit on Agilent 5300 Fragment Analyzer (Agilent Technologies, Palo Alto, CA, USA). RNA sequencing libraries were prepared using the NEBNext Ultra II RNA Library Prep Kit for Illumina following manufacturer's instructions (NEB, Ipswich, MA, USA). Briefly, mRNAs were first enriched with Oligo(dT) beads. Enriched mRNAs were fragmented for 15 minutes at 94 °C. First strand and second strand cDNAs were subsequently synthesized. cDNA fragments were end repaired and adenylated at 3'ends, and universal adapters were ligated to cDNA fragments, followed by index addition and library enrichment by limited-cycle PCR. Sequencing libraries were validated using NGS Kit on the Agilent 5300 Fragment Analyzer (Agilent Technologies, Palo Alto, CA, USA), and quantified by using Qubit 4.0 Fluorometer (Invitrogen, Carlsbad, CA). The sequencing libraries were multiplexed and loaded on the flowcell on the Illumina NovaSeq 6000 instrument according to manufacturer's instructions. The samples were sequenced using a 2x150 Pair-End (PE) configuration v1.5. Image analysis and base calling were conducted by the NovaSeq Control Software v1.7 on the NovaSeq instrument. Raw sequence data (.bcl files) generated from Illumina NovaSeq was converted into fastq files and de-multiplexed using Illumina bcl2fastq program version 2.20. One mismatch was allowed for index sequence identification.

After investigating the quality of the raw data, sequence reads were trimmed to remove possible adapter sequences and nucleotides with poor quality using Trimmomatic v.0.36. The trimmed reads were mapped to the Homo sapiens reference genome available on ENSEMBL using the STAR aligner v.2.5.2b. BAM files were generated as a result of this step. Unique gene hit counts were calculated by using feature Counts from the Subread package v.1.5.2. Only unique reads that fell within exon regions were counted. After extraction of gene hit counts, the gene hit counts table was used for downstream differential expression analysis. Using DESeq2, a comparison of gene expression between the groups of samples was performed. The Wald test was used to generate p-values and Log2 fold changes. Genes with adjusted p-values < 0.05 and absolute log2 fold changes > 1 were called as differentially expressed genes for each comparison. A gene ontology analysis was performed on the statistically significant set of genes by implementing the software GeneSCF. The goa_human GO list was used to cluster the set of genes based on their biological process and determine their statistical significance. A PCA analysis was performed using the "plotPCA" function within the DESeq2 R package. The top 500 genes, selected by highest row variance, were used to generate the plot.
